# Single cell atlas of the neonatal small intestine with necrotizing enterocolitis

**DOI:** 10.1101/2022.03.01.482508

**Authors:** Adi Egozi, Oluwabunmi Olaloye, Lael Werner, Tatiana Silva, Blake McCourt, Richard W. Pierce, Xiaojing An, Fujing Wang, Kong Chen, Jordan S. Pober, Dror Shoval, Shalev Itzkovitz, Liza Konnikova

**Affiliations:** Department of Molecular Cell Biology, Weizmann Institute of Science, Rehovot 7610001, Israel; Department of Pediatrics, Gynecology and Reproductive Sciences, Yale School of Medicine School, New Haven, CT 06520, USA; Program in Human and Translational Immunology, Gynecology and Reproductive Sciences, Yale School of Medicine School, New Haven, CT 06520, USA; Department of Obstetrics, Gynecology and Reproductive Sciences, Yale School of Medicine School, New Haven, CT 06520, USA; Pediatric Gastroenterology Unit, Edmond and Lily Safra Children’s Hospital Sheba Medical Center Ramat Gan 5262100, Israel; Sackler Faculty of Medicine, Tel Aviv University, Tel Aviv 6997801, Israel; Department of Medicine, University of Pittsburgh Medical Center Montefiore Hospital, Pittsburgh, PA 15213, USA; Department of Immunobiology, Yale School of Medicine, New Haven, CT 06520, USA

## Abstract

Necrotizing enterocolitis (NEC) is a gastrointestinal complication of premature infants with high rates of morbidity and mortality. A comprehensive view of the cellular changes and aberrant interactions that underlie this disease is lacking. Here, we combine single cell RNA sequencing, T Cell Receptor beta (TCRβ) analysis, bulk transcriptomics, and imaging to characterize cell identities, interactions and zonal changes in NEC. We find that inflammatory macrophages are abundant in NEC and that T cells exhibit increased expression of inflammatory genes and cytokines accompanied by an increase in TCRβ clonal expansion. Fibroblasts and endothelial cells increase in proportion and exhibit a switch to an activated pro-inflammatory state. Villus tip epithelial cell identity is substantially reduced in NEC and the remaining epithelial cells up-regulate pro-inflammatory genes. We establish a detailed map of aberrant epithelial-mesenchymal-immune interactions that may be driving inflammation in NEC mucosa. Our analyses highlight the cellular changes underlying NEC disease pathogenesis and identify potential targets for biomarker discovery and therapeutics.

## Introduction

Each year in the United States more than half a million infants are born prematurely. Necrotizing enterocolitis (NEC) is a devastating neonatal gastrointestinal complication associated with prematurity with high rates of mortality and morbidity. Its incidence is directly proportional to the degree of prematurity. Onset of symptoms occurs between 30-32 weeks corrected gestational age, irrespective of the gestational age at birth, or one to two months after delivery for the most premature infants (Yee et al. 2012). Current incidence of NEC is 1-7% of all the infants admitted to the Neonatal Intensive Care Unit with prevalence rising up to 15% for the most premature infants (Holman et al. 1997). A recent analysis of all-cause mortality in infants born prior to 29-weeks’ gestation showed that while overall mortality has declined, mortality related to NEC has increased (Patel et al. 2015). Moreover, in addition to short term complications, NEC is associated with high rates of long-term morbidity that include gastrointestinal (GI) strictures, feeding intolerance and short gut syndrome, but also significant systemic consequences such as microcephaly and neurodevelopmental delays (Neu and Walker 2011; Salhab et al. 2004).

NEC is a multifactorial disease involving environmental, microbial, host and immune factors. However, despite over one hundred and fifty years of research (Obladen 2009), the precise etiology of NEC continues to be elusive (Maheshwari et al. 2014), without effective prevention methods or treatment options available.

Although dysregulation in both the mucosal immune system and the epithelial barrier are hypothesized to be associated with NEC, the mechanism of how these cells contribute to disease onset or progression is not clear. Previous studies have typically focused on specific cellular populations individually (Denning et al. 2017; Hackam et al. 2013) and a comprehensive unbiased analysis is lacking. To thoroughly address these questions, studies at the single cell level utilizing primary human tissue are needed. Emerging data suggest that fetal intestinal mucosal immunity is established by as early as 14 weeks’ gestation with phenotypical tissue resident memory (TRM) T cells residing in the small intestine (SI) (Stras et al. 2019; Schreurs et al. 2019; Li et al. 2019). These data allude that the etiology of increased susceptibility to infection and therefore NEC in preterm infants is unlikely to be simply due to immaturity but rather caused by underlying defects in immune function. As such, an understanding of the neonatal SI at a system’s biology level is needed to further our understanding of NEC pathogenesis.

With the goal of developing better treatments for premature infants with NEC, here we reconstruct an atlas of SI neonatal and NEC tissue with associated cellular localizations and interactions. Using single cell RNA sequencing (scRNAseq) data we define transcriptional signatures and ligand-receptor interactions associated with NEC. This is further combined with deconvoluted bulk RNAseq, mass cytometry and imaging analysis to define cellular proportions, locations, and interactions. Our data demonstrate that aberrant epithelial-mesenchymal-vascular-immune cell interactions contribute to NEC associated intestinal inflammation. Specifically, we describe an increase in inflammatory macrophages and inflammatory changes in conventional and regulatory T cells accompanied by increased T Cell Receptor beta (TCRβ) clonality. This is further correlated with profound remodeling of NEC mucosa with proportional decrease in top villus epithelial cells and increase in endothelial and fibroblast cells, where all three cell types increase transcription of inflammatory genes. Our NEC single cell atlas identifies networks controlling intestinal homeostasis and inflammation bridging the gap in the incomplete understanding of NEC pathogenesis and biomarker and therapeutics discovery.

## Results

### A single cell atlas of NEC and control human small intestinal samples

To characterize the cell states associated with NEC we extracted SI tissues from NEC patients and from neonatal samples without NEC (**Fig 1a**). We performed scRNAseq on 11 subjects (6 NEC and 5 neonatal samples **Fig 1b-c**). Since estimation of cell abundances in single cell atlases can be skewed due to differential viability of extracted cells, we additionally performed bulk RNAseq computational deconvolution (10 NEC and 4 neonatal samples), as well as suspention and imaging mass cytometry (CyTOF:12 NEC and 6 neonatal samples; IMC: 5 NEC and 3 neonatal samples, **Fig 1a**). We also implemented next generation sequencing (NGS) of TCRβ to identify clonality changes (5 NEC and 9 neonatal samples). Furthermore, we combined scRNAseq atlas and IMC data to define niche and cellular interactions enriched in NEC (**Fig 1a**).

**Figure 1.**
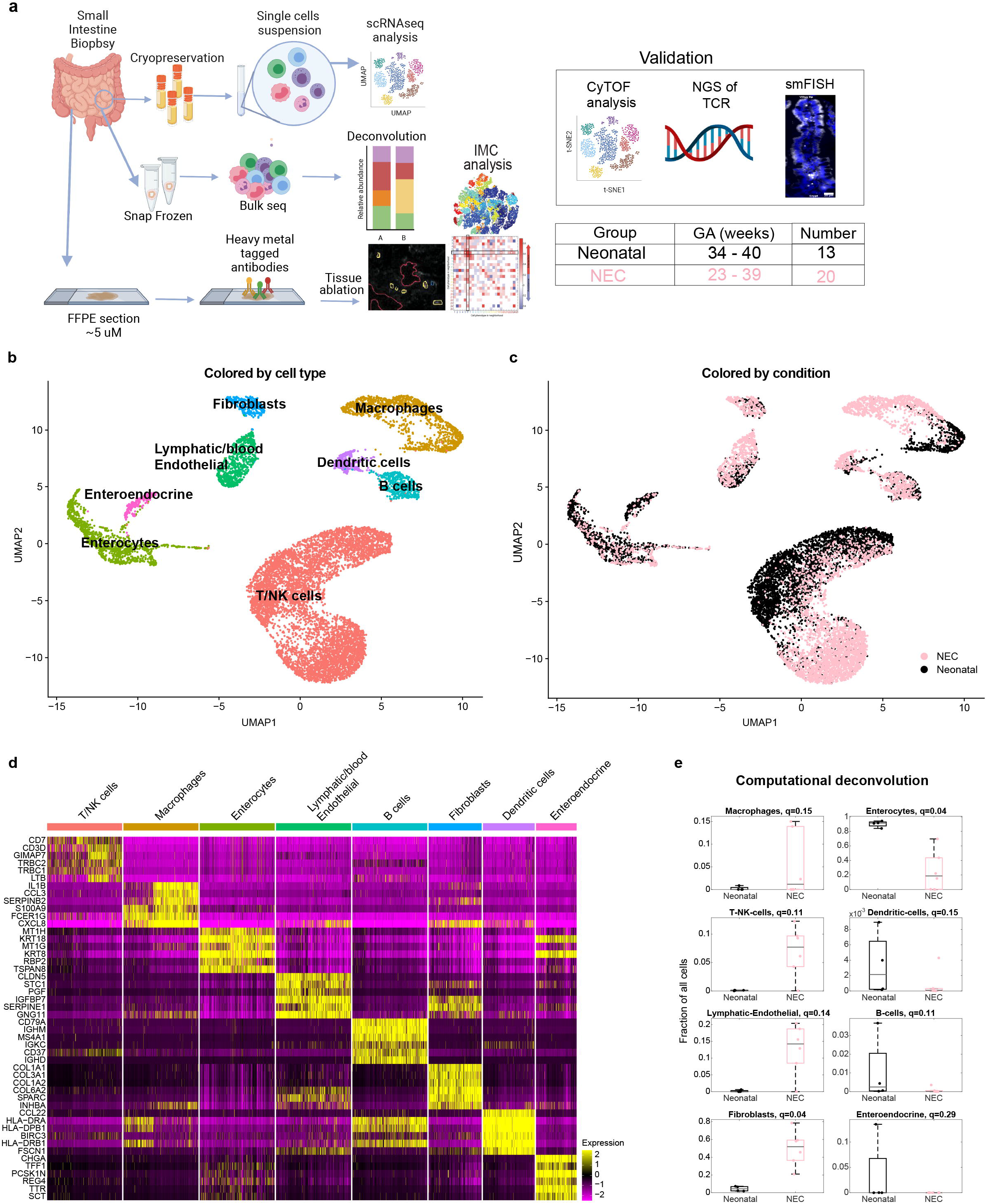
A single cell atlas of NEC and control human small intestinal samples. **a** – Experimental layout – human small intestinal tissues from neonates and NEC patients were harvested and used for single cell RNAseq, bulk RNAseq, smFISH, imaging mass cytometry (IMC) and suspension mass cytometry (CyTOF). **b** – Single cell atlas annotated by cell type. **c** – Single cell atlas annotated by condition. **d** – Markers of the cell types in **b. e-** Estimates of the proportion of T-NK cells, Macrophages, enterocytes, lymphatic/blood endothelial cells, fibroblasts, dendritic cells and enteroendocrine cells based on computational deconvolution of the bulk RNAseq using the atlas single cell populations (Methods). Each dot is a sample, fractions normalized to the sum of cell fractions; q-values are computed based on FDR correction for all cell populations in the full atlas (Methods). Gray lines are medians, black/pink boxes are 25-75 percentiles.

Our atlas included 11,308 cells and revealed 8 main cell clusters representing macrophages, dendritic cells, B cells, T/NK cells, lymphatic/blood endothelial cells, fibroblasts, enteroendocrine and enterocytes populations (**Fig 1b-d**). Each cluster exhibited distinct gene expression markers (**Fig 1d, Supplementary Table 1**). Our atlas enabled exploring the detailed gene expression changes in distinct cell subsets. Furthermore, bulkseq analysis followed by computational deconvolution facilitated determination of major cluster abundances (**Fig 1e**). This analysis revealed presence of inflammatory macrophages, dendritic cells (DCs) and T/Natural Killer (NK) cells with an aincrease in lymphatic/endothelial cells and fibroblasts and a decrease in enterocytes.

To complement the bulk RNAseq analysis in assessing cell-type abundances we applied IMC. In this method, formalin-fixed paraffin-embedded sections of small intestine are incubated with a cocktail of antibodies (**Supplementary Table 2)**, ablated and analyzed. Clustering analysis of 20,819 cells from IMC data revealed numerous clusters inclusing epithelial cells, endothelial cells, fibroblasts, B cells, T cells, macrophages, DCs and clusters of cells that could not be identified with the markers used (other, **Fig S1a-b**). The IMC data facilitated acquisition of cellular abundance and niche cell interactions in the native state. Our panel was also designed to identify signaling cascades by incorporating antibodies against phosphorylated proteins.

### NEC is associated with an increase in inflammatory genes in macrophages and dendritic cells

Myeloid cells have been shown to be enriched in NEC tissue and to exhibit elevation in inflammatory programs (Zhang et al. 2021; Olaloye et al. 2021; Xia et al. 2021; Liu et al. 2019). Our atlas included 1,836 myeloid cells, which clustered into dendritic cells, non-inflammatory macrophages, as well as two separate clusters of inflammatory macrophages (**Fig 2a-c**). One of the inflammatory macrophage clusters, made up almost exclusively of NEC associated macrophages (Inflammatory macrophages group A), showed elevation of inflammatory cytokines and chemokines such as *IL6, IL1B* and *CXCL8* (**Fig 2c**). Accordingly, differential gene expression between the NEC and neonatal macrophages showed a significant increase in pro-inflammatory molecules, including *IL1A, IL1B, IL6, CSF2, CSF3, CXCL8, CCL3, CCL4, CXCL2, CCL20* and signaling molecules such as *IRAK2, SOD2, NFKBIz, and NFKBIA* (**Fig 2c-d, Methods**). NEC macrophages exhibited downregulation of anti-inflammatory genes such as *CD9, MERTK, HLA-DRB5, TGFBI*, and *PLXDC2* (Gong et al. 2012, **Fig 2d**). Similarly, dendritic cells in NEC up-regluated genes associated with inflammation such as *ISG15, S100A6, SERPINB1, GNG11, CCL17* and *FSCN1* (Dantoft et al. 2017) and genes involved in activation - *CD40* and *CD44* (Yeh et al. 2021; Buhtoiarov et al. 2005) (**Fig 2e, Supplementary Table 3**). Macrophages were enriched in inflammatory gene sets, such as TNF*α* signalling via NF-*κ*B, inflammatory response, NOD like receptor (NLR) signaling and Toll Like Receptor (TLR) signaling (**Fig 2f**). Both NLR and TLR activation have previously been implicated in NEC pathogenesis and NF-*κ*B signaling in inflammation in general (Afrazi et al. 2014; Hackam and Sodhi 2018; Cushing et al. 2017). Dendritic cells in NEC were enriched in MYC target genes and ribosomal pathways (**Fig S2a**). Myc signaling in DCs has been shown to be critical for optimal T cell priming (Kc et al. 2014). Computational deconvolution of the bulk RNAseq samples revealed a trend toward an increase in overall macrophages (fold change=67, q=0.15, **Fig 1e**) with an increase in the inflammatory macrophages (fold change = 667, q=0.02, **Fig 2g**) and a decrease in the non-inflammatory macrophages in the NEC samples (fold change = 0.009, q=0.02, **Fig 2g**), while dendritic cells were overall decreased in NEC (fold change =0.056, q=0.29, **Fig 2g**). Using CyTOF, we confirmed an increase in the overall macrophages (fold change=1.46, p=0.09, **Fig 2h**) and in the inflammatory macrophages (CD16^+^, fold change=1.32, p=0.05, **Fig 2h**). We also noted a trend towards a decrease in dendritic cells in particular the CD103^+^ and CCR7^+^CD103^+^ subtypes, known to be the canonical antigen presenting cells that traffic to the mesenteric lymph nodes, in NEC comparing to neonatal samples (fold change=0.35, p=0.07 **Fig 2h, S2b**).

**Figure 2.**
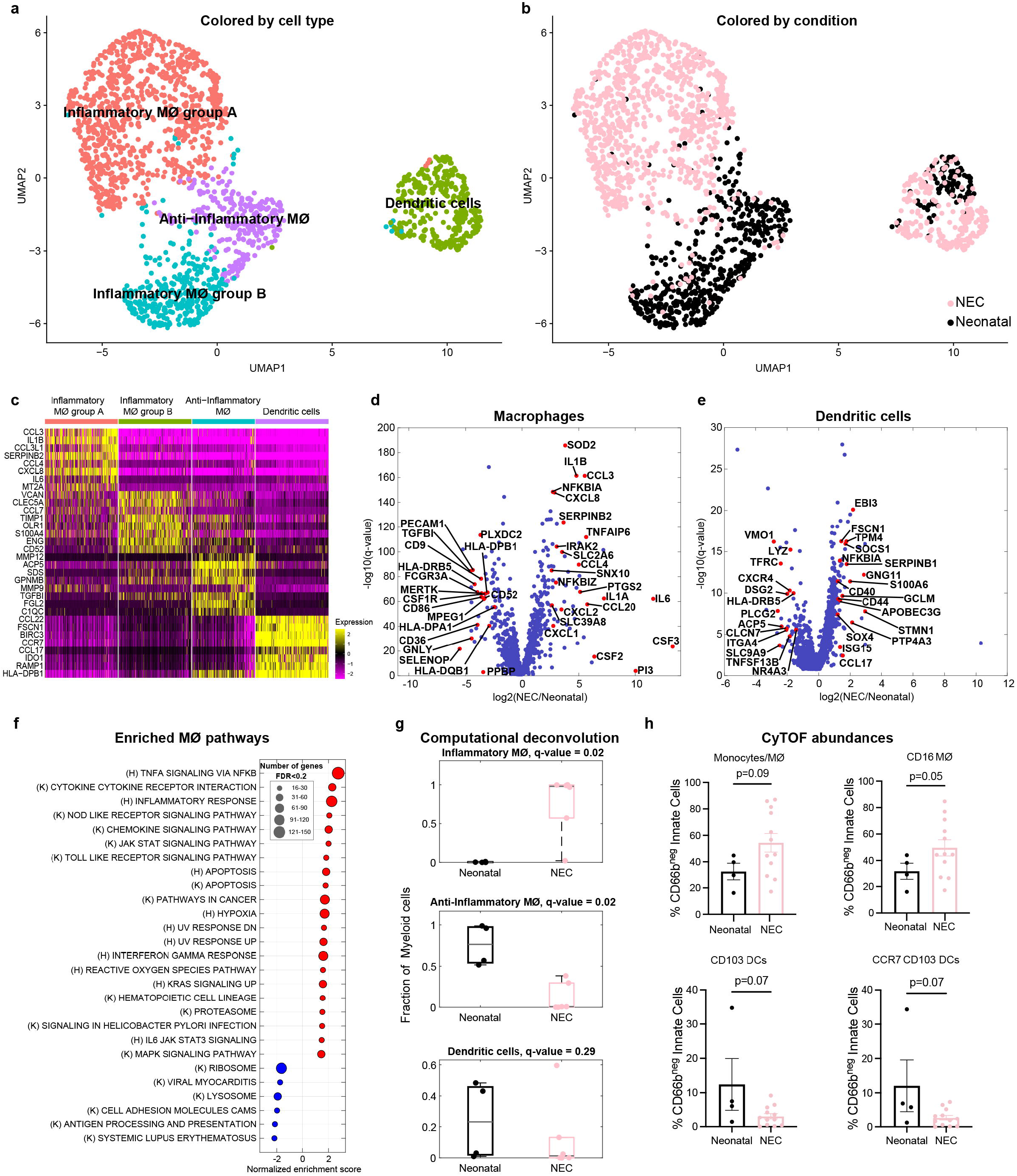
Inflammatory macrophages are increased in NEC. **a** – Re-clustered atlas of myeloid lineages. Mϕ– macrophages. **b** – Single cell atlas annotated by condition. **c** – Top 8 markers of the myeloid cell sub-types. **d** – Differential gene expression between NEC and neonatal macrophages. **e** – Differential gene expression between NEC and neonatal dendritic cells. Red dots (d,e) are selected differentially expressed genes among the genes with q value<0.02 and fold change above 3 or below 1/3. Included are all genes with normalized expression above 10^−4^. **f** – Gene set enrichment analysis (GSEA) of pathways enriched (red) or depleted (blue) in NEC samples compared to neonatal samples for macrophages (q value<0.2). **g** – Estimates of the proportions of distinct myeloid cell subsets, based on computational deconvolution of bulk sequencing data. Each dot is a sample, proportions were renormalized over all myeloid cells, q-values are computed based on FDR correction for myeloid cells only (Methods). Gray lines are medians, black/pink boxes are 25-75 percentiles. **h** – Monocytes/Mϕ and dendritic abundances in neonatal and NEC samples from mass cytometry data adapted from (Olaloye et al. 2021) represents abundances from Fig S2b.

### T cell numbers, clonality, and characteristics are changed in NEC

The contribution of lymphocytes to the pathogenesis or progression of NEC has been controversial, with some studies indicating an increase in T cells while others showing a decrease (Weitkamp et al. 2014; Anttila et al. 2003; Weitkamp et al. 2013). Moreover, work from the Hackam group has suggested that Th17 lymphocytes are critical to NEC pathogenesis (Egan et al. 2016; Jia et al. 2019). Our atlas included 6,400 T/NK cells, with 7 distinct T and NK cell subsets inculding: NK cells, innate lymphoid cells (ILCs), naïve T cells, regulatory T cells (Tregs), tissue resident memory (TRM) T cells, proliferating T cells and activated T cells (**Fig 3a-b, S3a**).

**Figure 3.**
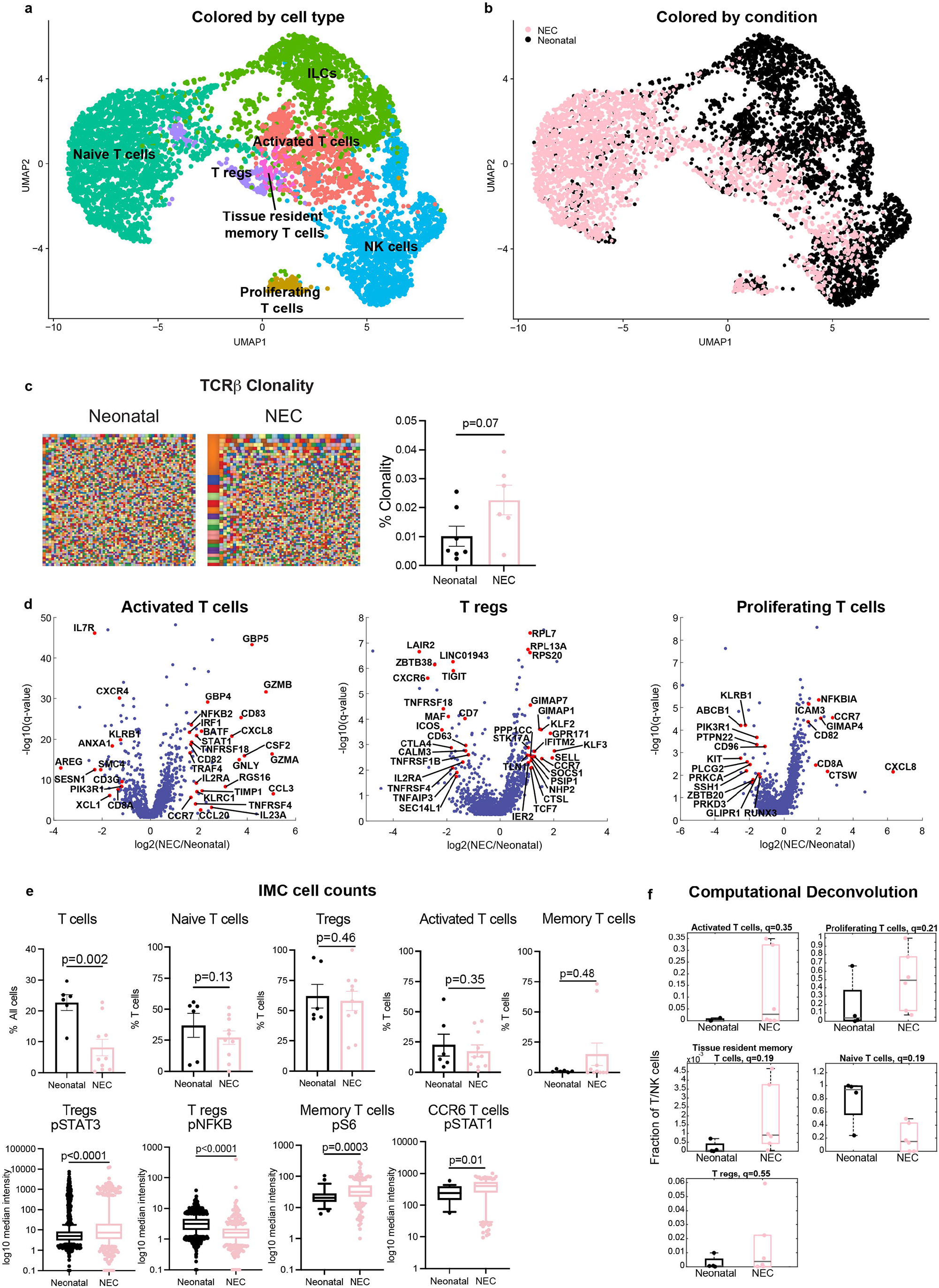
Landscape and transcriptional signatures of T/NK/ILC populations in NEC. **a** – Re-clustered atlas of the T/NK cluster. **b** – Single cell atlas annotated by condition. **c** – Next generation sequencing of T cell receptor β (TCRβ) candy plots where each small square represents one clone with the squares proportional to the number of T cells with a particular clone with quantification on the right. **d** – Differential gene expression between NEC and neonatal T cell populations. Red dots are selected differentially expressed genes among the genes with q value<0.02 and fold change above 2 or below 1/2. Included are all genes with normalized expression above 10^−4^. **e-** Cell counts of T cells from imaging mass cytometry (IMC) images, where each dot represents 1 image (Methods). Abundance of naïve T cells, T regs, activated T cells and memory T cell clusters from Fig S1a-b expressed as percentage of T cells. Boxplot of Log10 of mean metal intensity for individual cells. Expression of phosphorylated STAT3 (pSTAT3) and pNFKB on T regs, phosphorylated ribosomal protein S6 (pS6) on memory T cells, and phosphorylated STAT1 (pSTAT1) on CCR6 T cells. Each dot represents one image. Two images scanned per sample (Neonatal, n=3, NEC, n=5). P-values computed using Wilcoxon rank-sum test. (Methods). Boxes show 25-75 percentiles, with whiskers extend to to 10-90 percentiles. **f-** Estimates of the proportions of activated T cells, proliferating T cells, tissue resident memory (TRM) T cells, naïve T cells and Tregs based on computational deconvolution of the bulk sequencing data, using the atlas single cell populations (Methods). Each dot is a sample, proportions were renormalized over all T cells, q values are computed based on FDR correction for T cells subsets only.

Given the abundance of memory T cells in the fetal SI (Schreurs et al. 2019; Stras et al. 2019) and their presence in neonatal and NEC tissue (**Fig 3**), we sought to determine if there was a clonal expansion of T cell populations in NEC using NGS of the TCRβ. TCRβ clonality was increased in NEC compared to neonatal samples (**Fig 3c**). Additionally, analysis of gene usage revealed that the use of variable (V), diversity (D) and joining (J) segments differed between NEC and neonatal cases with an increased frequency of TRBV10 and decreased use of TRBV15, TRBJ1-4 and TRBJ2-1 in NEC compared to neonatal cases (**Fig S3b-c)**. NEC patients had shorter CDR3β length with fewer deletions and a trend towards fewer insertions than neonatal controls (**Fig S3d)**, a phenomenon previously reported in IBD (Werner et al. 2019).

To determine which clones were expanded in NEC, we performed a search of the public clones database. Public clones have a unique amino acid or nucleotide sequence, are present across individuals and can be easily identified in a published database (https://vdjdb.cdr3.net/). The majority of the top clones observed in NEC were previously seen public clones. Intrestingly, the amino acid sequence of three clones enriched in NEC had an identical sequence to clones known to bind to cytomegalovirus (CMV) (CMV-1, CMV-2, CMV-3) (**Fig S3e, Supplementary Table 4**).

To further investigate if T cells, ILCs and NK cells were transciptionally altered in NEC, we compared the transcriptomes of all T cell clusters between NEC and neonatal samples. Differential gene expression (DGE) showed an increase in an inflammatory signature in all subtypes except the naïve T cells (**Fig S3f**). No significant alterations in the abundance of ILCS or NK cells were observed in the deconvolution analysis (Fig S3g). The ILC cluster had an increase in genes associated with trafficking, *CCR7* (**Fig S3**h) while the NK cluster had an increase in *CCL3* (Marischen et al. 2018), *CD83, XCL, GZM* and TNF-related genes (**Fig S3i**). Similarly, the activated T cell cluster showed an upregulation of inflammatory and cytotoxic genes such as *GZMA/B, GNLY, KLRC1, CSF2, IL23A, CXCL8*; genes indicative of activation such as *IL2RA*, and genes downstream of IFN-*γ* signaling such as *IRF1, GBP5* and *STAT1* (Ivashkiv and Donlin 2014) (**Fig 3d**). We could not detect *IL17A* expression and *IL17F* and *IL22*, also produced by TH17 cells, were only upregulated in 1 NEC case and did not meet the threshold to be included in the DGE analysis. However, a number of other IL17 signature genes were upregulated in activated T cells assoicated with NEC including *CCL20, TIMP1* (also plays a role in TH1 cells) and *BATF* (Kurachi et al. 2014) (**Fig 3d**). We could not perform DGE for the TRM cluster as there were very few neonatal T cells contributing to it. Yet, IMC analysis of memory T cells showed an increase in pS6 indicative of mTOR signaling associated with increased glycolytic activity associated with effector function (**Fig 3e** (Salmond et al. 2015)).

### Regulatory T cells with reduced immunosuppressive signature associated with NEC

Tregs have an important role in regulating mucosal homeostasis. Conflicting reports have suggested their contribution to NEC (Pang et al. 2018a; Weitkamp et al. 2013; Dingle et al. 2013). Using IMC, we found an overall reduction in T cell numbers (fold change=0.3, p=0.002, **Fig 3e**, a non-significant reduction in naïve T cells (fold change=0.1, p= 0.13 **Fig 3e)** and no difference in the proportion of Tregs (fold change 1.26, p=0.46, **Fig 3e)**. Computational deconvolution revealed a decrease in naïve T cells, (fold change = 0.16, q=0.19, **Fig 3f**), an increase in TRM T cells (fold change =19, q=0.19, **Fig 3f**) and a trend towards an increase in activated T cells (fold change = 26, q=0.35, **Fig 3f**), without a change in Tregs (**Fig 3f**). Analysis of DGE in Tregs from the scRNAseq data revealed an increase in *SELL* (CD62L), *CCR7, SOCS1* and a number of ribosomal genes, potentially indicating increased translation (**Fig 3d, Supplementary Table 3**). Interestingly, there was a reduction in the expression of a number of genes classically associated with Treg identity or supression function such as *CTLA-4, TIGIT, iCOS, TNFRSF4* (Ox40-found on intestinal Tregs and effector memory T cells) and *IL2RA* (**Fig 3d, Supplementary Table 3**). IMC analysis demonstrated an increase in pSTAT3 and a decrease in pNFkB signaling in NEC associated Tregs (**Fig 3e**). In summary, our analysis indicates that Tregs do not significantly change in amount in NEC, but rather show features of reduced suppressive activity.

### NEC blood and lymphatic endothelial cells exhibit distinct pro-inflammatory signatures

Our single cell atlas included 407 vascular endothelial and 332 lymphatic endothelial cells, each clustering into pro-inflammatory and non-inflammatory subsets (**Fig 4a-c**). *CLDN5* marked all endothelial cells and *PDPN* was specific to the lymphatic endothelial clusters (**Fig S4a-b)**. Upon examination of the lymphatic marker PDPN, we observed that the lymphatic endothelial cell cluster contained a subset that were potentially not lymphatic but rather blood endothelial cells (**Fig S4b, red star**). However, there was no difference in DGE or pathway analyses if those few cells were removed. Additionally, we observed CCL21 expression restricted to lymphatic endothelial cells, minimal CCL19 expression in any neonatal endothelial cells and CXCL10 expression in inflamatory lymphatic endothelial cells (**Fig S4c**).

**Figure 4.**
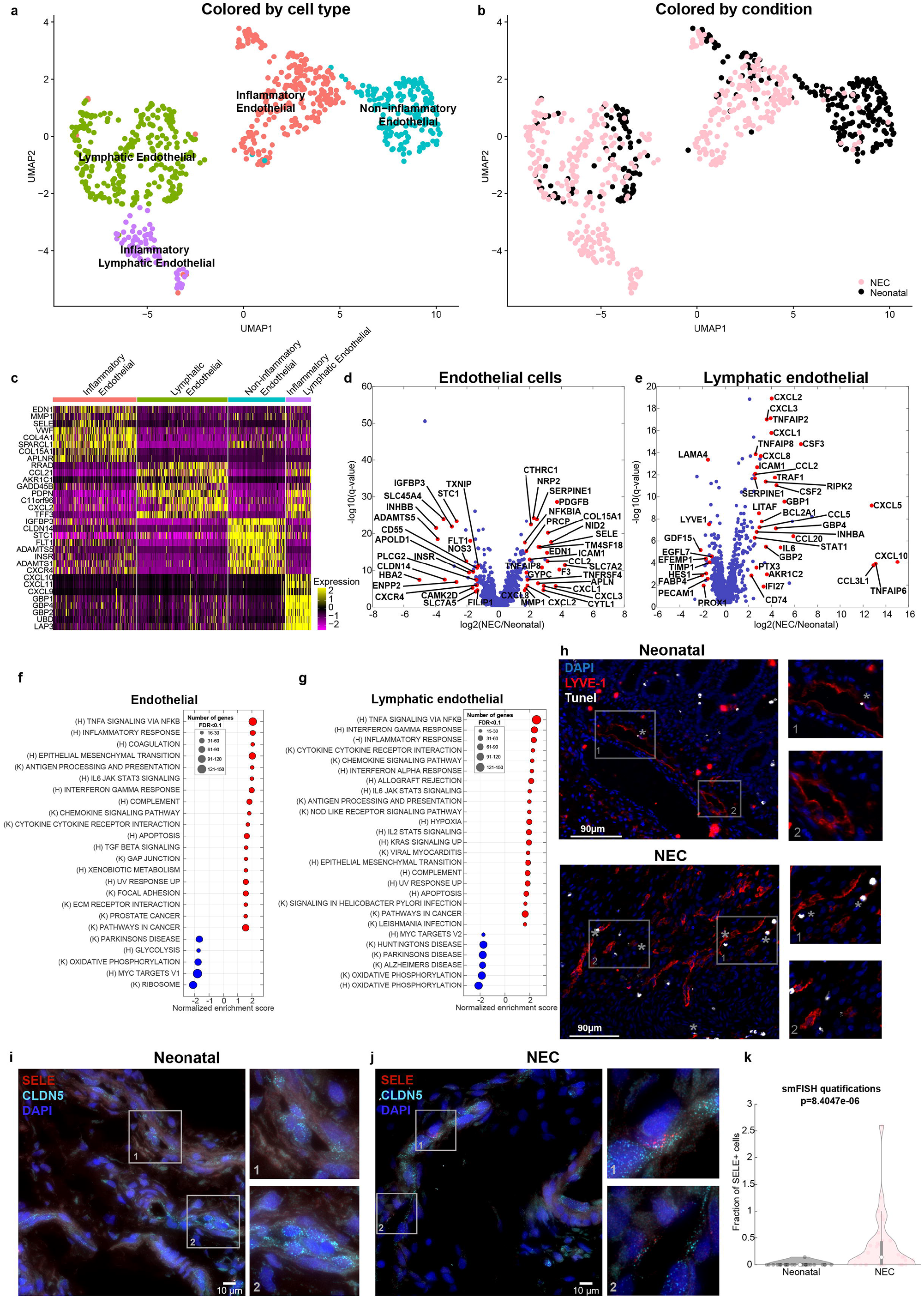
Lymphatic and blood endothelial cells in NEC exhibit pro-inflammatory signatures. **a** – Re-clustered atlas of the lymphatic and blood endothelial cluster. **b** – Single cell atlas annotated by condition. **c** –Markers for the cell types in **a. d-e** – Differential gene expression between NEC and neonatal cells for blood endothelial cells (d) and lymphatic endothelial cells (e). Included are all genes with normalized expression above 10^−4^. Red genes are selected differentially expressed genes among the genes with q value<0.02 and fold change above 2 or below 1/2. **f-g** – GSEA of pathways enriched (red) or depleted (blue) in NEC samples compared to neonatal samples for blood endothelial (f) and lymphatic endothelial cells (g) with q value<0.1. **h** - Representative immunofluorescence images from neonatal and NEC samples stained with Lyve-1(red) and TUNEL staining (white). Grey asterisk (*) represent apoptotic endothelial cells (Tunel^+^LYVE-1^+^). **j-l** – smFISH demonstrating increase in SELE^+^ endothelial cells in NEC. Red dots are individual mRNAs of SELE, cyan dots are individual mRNAs of CLDN5, a marker of blood/lymphatic endothelial cells. Blue are DAPI-stained nuclei, scale bar – 10µm. **l–** Quantification of smFISH signals of SELE^+^ cells in NEC (n=2) compared to a neonatal sample (n=2). Each dot represents the number of SELE+ cells in one imaging field.

Upon DGE analysis, we found that NEC blood and lymphatic endothelial cells uniformly upregulated chemokines such as *CCL2, CXCL1, CXCL2* and *CXCL3* as well as adhesion molecule, *ICAM1*, consistent with endothelial activation (Pierce et al. 2017, **Fig4d-e**). In blood endothelial cells, we observed upregulation of adhesion molecules *SELE* (Golias et al. 2011), procoagulant factors such as *SERPINE1* and *F3* along with changes in regulators of blood flow that reduce perfusion including increased *EDN1* and decreased *NOS*3 (**Fig 4d)**. We observed significant upregulation of cytokines and chemokines in lymphatic endothelial cells including *IL6, CCL5, CXCL10* (Hillyer et al. 2003), *CSF2, CSF3, CCL20, CXCL8* and pro-inflammatory signaling such as *TNFAIP2, TNFAIP6, TNFAIP8, TRAF1* (where binding to TNFR2 enhances CXCL8 production (Wicovsky et al. 2009)) and *RIPK2* (**Fig4e**). Pathway analysis revealed both blood and lymphatic endothelial cells were significantly enriched in NFκB-responsive genes, chemokine signaling pathway and IFN*γ* responses (**Fig 4f-g**). Both *IL1B* and *TNFA* were enriched in the NEC associated inflammatory Mϕ that correlated with the increase in *SELE* and *ICAM1* in the activated endothelium in NEC (**Fig S5a-d)**. Apoptotic gene sets were also enriched in both blood and lymphatic endothelial cells (**Fig 4f-g**), confirmed by an increase in TUNEL staining in LYVE-1^+^ cells (**Fig 4h, S4d**). To validate the induction of pro-inflammatory endothelial cells in NEC, we performed single molecule fluorescence in-situ hybridization (smFISH) for the endothelial/lymphatic specific marker *CLDN5* along with *SELE* as a specific marker of activation endothelial activation (**Fig 4i-j**). We found a significantly higher abundance of SELE^+^ endothelial cells in the NEC samples (0 vs. 2, p=2.5×10^−9^, **Fig 4k**).

Deconvolution of bulkseq revealed an overall increase in blood/lymphatic endothelial cells (fold change= 833, q=0.14, **Fig 1e**) with a non-significant increase in endothelial cells (bulk RNAseq fold increase = 2.7 (inflammatory endothelial cells), 1.8 (endothelial cells), q=0.38, 0.67, **S5e)**. Similarly, immunoflorescence for LYVE-1 staining showed an increase in NEC (fold change=1.6, p=0.02, **Fig S4d**). In summary, we found a proportional increase in endothelial cells in NEC with a shift in blood and lymphatic endothelial cells towards a pro-inflammatory state complemented by increased coagulation and decreased perfusion-associated genes in the vascular endothelium.

### NEC epithelial cells and fibroblasts exhibit an increased inflammatory potential with a loss of villus-tip enterocytes

We next turned to analyze the changes in enterocyte cell identity in NEC. Our atlas included 1,203 enterocytes (**Fig 5a**-**b**), which we classified by crypt-villus zones using known landmark genes (**Fig 5c-e**). Computational deconvolution exposed an overall reduction in enterocytes (fold change= 0.2, q=0.04, **Fig 1e**) with a substantial reduction in villus top cells (fold change =0.3, q=0.2, **Fig 5f**) and an increase in crypt cells in NEC compared to neonatal (fold change = 76, q=0.2 **Fig 5f**). IMC data also demonstrated a reduction in epithelial cells (**Fig 5g**) and vilous bluntng (**Fig 5h**). DGE of the lower villus and crypt zones revealed that NEC epithelial cells exhibited substantial increase in the anti-microbial genes *LCN2, REG1A, REG1B* and *DMBT1*, genes previously shown to be elevated in inflammed epithelium associated with inflammatory bowel disease (IBD) (van Beelen Granlund et al. 2013; Kaemmerer et al. 2012). NEC enterocytes further increased the expression of the chemokines *CXCL1, CXCL2, CXCL3* (Puleston et al. 2005), *CXCL5* and *CXCL8* (**Fig 5i, Supplementary Table 3**). Likewise expression of *DUOXA2*, an IBD-associated gene that produces H_2_O_2_ (MacFie et al. 2014) was increased in NEC epithelial cells. Additonally, signaling in epithelial cells was altered in NEC with a notable increase in pSTAT1, pSTAT3, and pS6 (**Fig 5g**). This is consistent with increased expression of STAT3 target genes: *REG1A, REG1B, LCN2, DMBT1, CXCL5* and STAT1 target gene *DUOXA2* (Wu et al. 2011).

**Figure 5.**
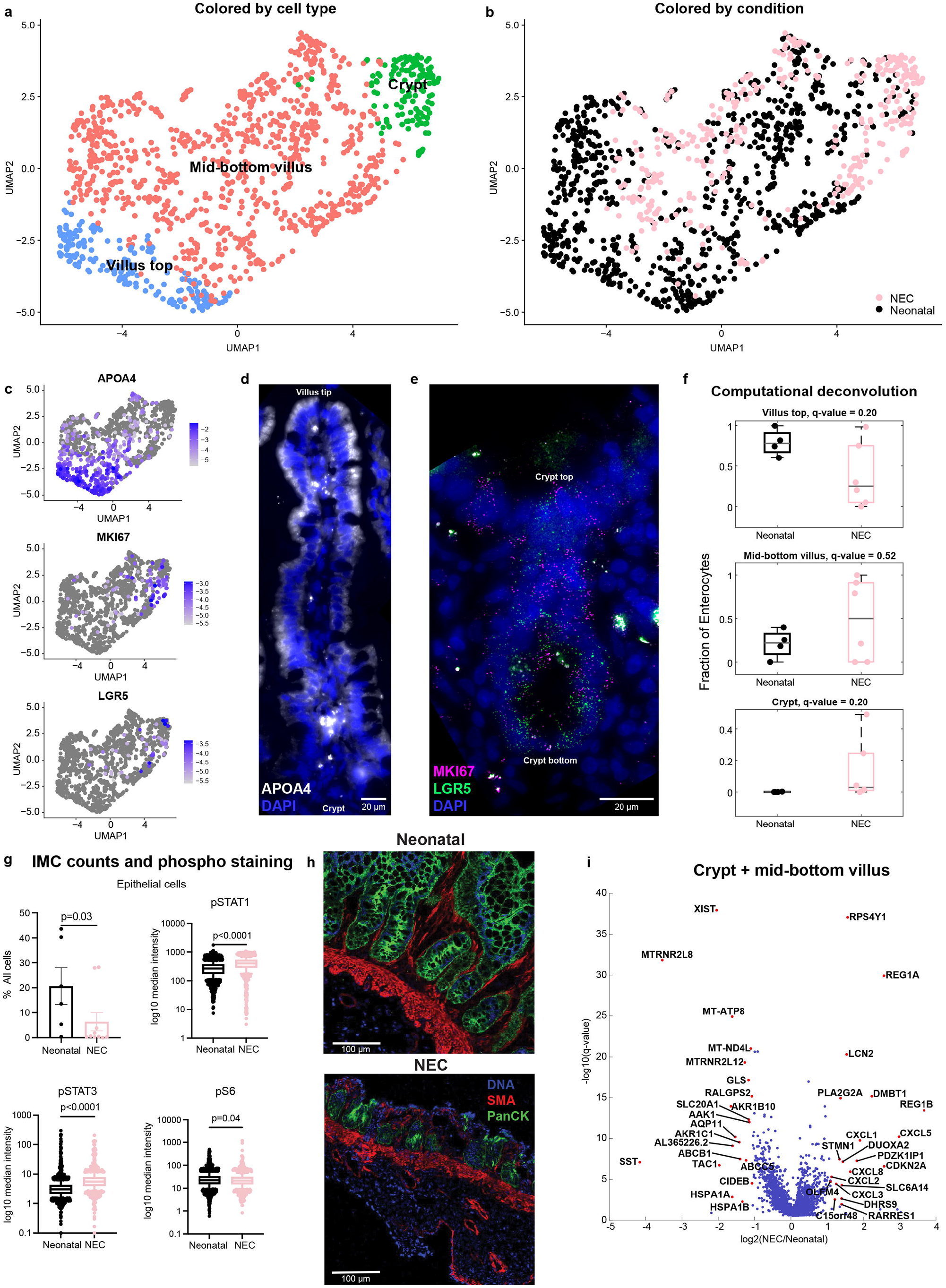
Changes in enterocyte gene expression and zonal representation in NEC. **a** – Re-clustered atlas of the enterocyte cluster colored by crypt-villus zone. **b** – Single cell atlas annotated by condition. **c** – UMAPs colored by top villus marker - APOA4, proliferation marker – MKI67 and stem cell marker – LGR5. Color bar is log10 (normalized expression). **d-e** – smFISH of epithelial cells (d) demonstrates increase in APOA4^+^ (white) epithelial cells towards the top of the villus. **e** – Magenta dots are individual mRNAs of MKI67, green dots are individual mRNAs of LGR5 in the crypts. Blue are DAPI-stained nuclei, scale bar – 20µm. **f** – Estimates of the proportions of villus-crypt zones subsets, based on computational deconvolution of bulk sequencing data. Each dot is a sample, proportions were renormalized over all villus-crypt zones, q-values are computed based on FDR correction for enterocytes only (Methods). **g** – IMC quantification of epithelial cells and median log10 mean metal intensity of phosphorylated signaling markers for mTOR pathway (pS6), and STAT pathways (pSTAT3 and pSTAT1). **h –** Representative images from Histocat® 1.7.6.1 showing villus blunting in NEC compared to neonatal tissue. DNA – 191/193-intercolator (blue), SMA-smooth muscle actin (red), panCK-pancytokeratin (green). **i-**Differential gene expression between NEC and neonatal cells for the crypt and mid-bottom villus zones. Included are all genes with normalized expression above 5×10^−5^. Red dots are the top 20 most differentially expressed genes among the genes with q value<0.02 and fold change above 2 or below 1/2.

Clustering of the fibroblasts demonstrated a small neuronal population and 319 fibroblasts (**Fig S6a-b**). Using computational deconvolution, we identified an increase in the overall fibroblasts (fold change=11.7, q=0.04, **Fig 1e**). NEC fibroblasts exhibited an increase in several inflammatory genes including *IL1B, CSF2, CSF3, EREG* and *CCL20* (**Fig S6c, Supplementary Table 3**).

### Epithelial-mesenchymal-immune interactions are altered in NEC

Our analysis identified a massive infiltration of inflammatory macrophages, alterations in endothelial and epithelial cells and a shift towards an inflammatory phenotype in lymphocytes. We next asked what are the niche signals that could be associated with elevated recruitment of these inflammatory immune cells. To define the cellular interactions in NEC SI we performed a nearest neighbor analysis of IMC data using histoCAT, a tool that identifies statistically significant interactions/avoidances between cellular clusters (Schapiro et al. 2017). Numerous cell type interactions were altered in NEC (**Fig 6a, Supplementary Table 5**). Consistent with upregulation of genes associated with leukocyte recruitment on endothelial cells such as SELE (**Fig 4d-g, S5b**,**d**), NEC tissue had increased interactions of endothelial cells with DCs. Additionally, fibroblasts had increased interactions with epithelial cells, with monocytes and crypt epithelial cells. Epithelial cells had increased interactions with monocytes. Consistent with increase in T cell clonality, NEC SI showed significant interactions between memory T cells and antigen presenting cells including DCs and macrophages. Finally, Tregs had significantly altered interactions with several other cell types including decreased interactions with macrophages (**Fig 6a**).

**Figure 6.**
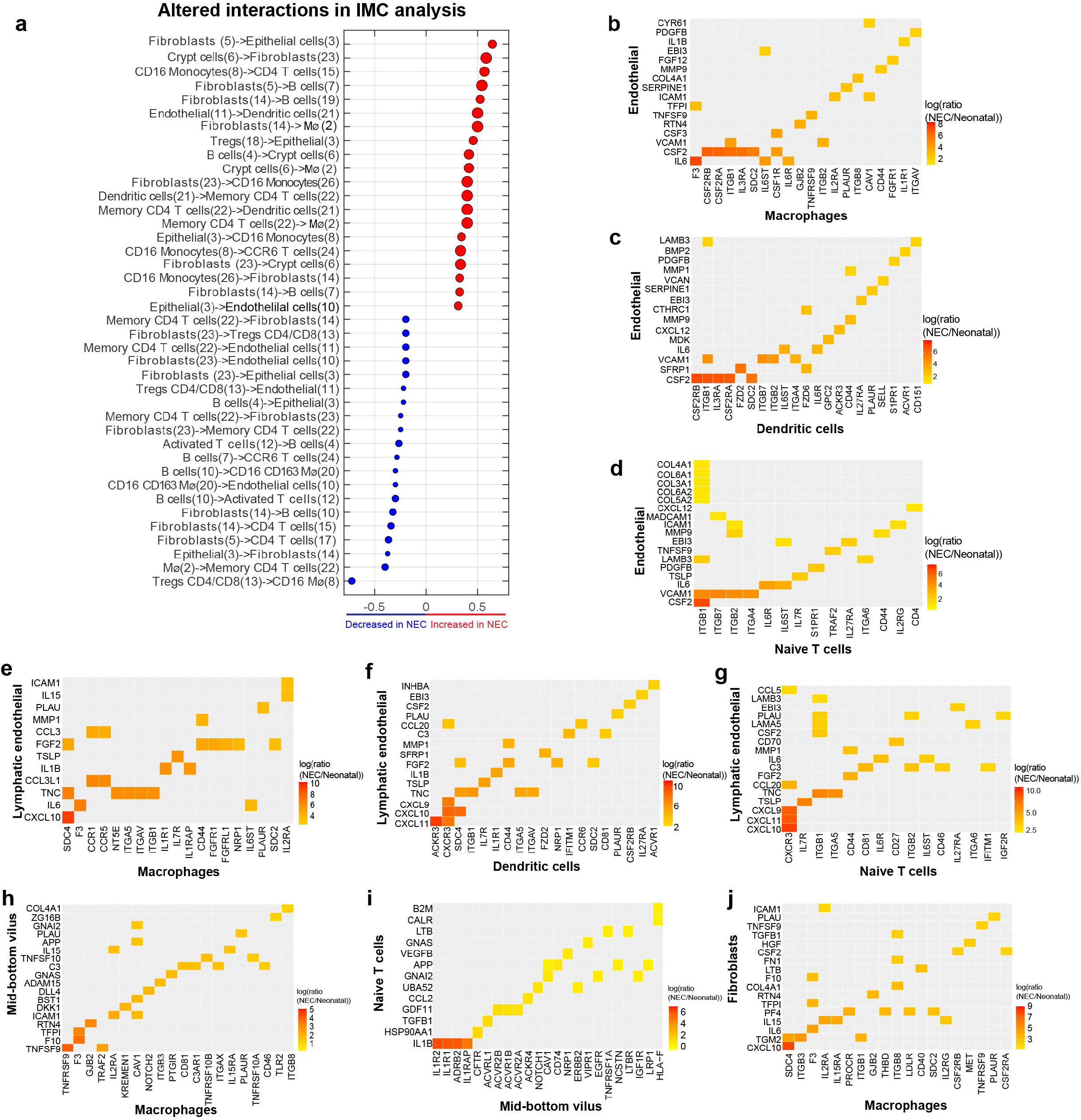
Altered cellular adjacencies and protein-ligand interactions in NEC **a**- Dot plot showing the 20 interaction types that have the highest increase (red) or decrease (blue) interaction values between neonatal and NEC samples (Methods). Interactions values (**Supplementary Table 5)** were computed by imaging mass cytometry analysis using Histocat 1.7.6.1 and 999 permutations and a p value <0.01 (Schapiro et al. 2017). Dot size corresponds to the interaction values in NEC. **b-d** Significantly elevated molecular interactions between blood endothelial cells and macrophages (b) dendritic cells (c), naïve T cells (d). **e-g** Significantly elevated interactions between lymphatic endothelial cells and macrophages (e), dendritic cells (f) and naive T cells (g). **h-j** -Interactions between epithelial cells and macrophages (h) naïve T cells and epithelial cells (i) or fibroblasts (j) and macrophages. Shown are 16-25 significant interactions (q value<0.01) with highest fold change (Methods). In all interaction maps sender population is on the y axis, received is on the x axis.

To identify the chemokines and cytokines that could be associated with recruitment of inflammatory macrophages and lymphocytes or could be responsible for altered signaling within the NEC SI, we performed a ligand-receptor analysis between all pairs of cell types in our atlas (**Supplementary Table 6, Methods**). To this end, we parsed a database of ligands and matching receptors (Ramilowski et al. 2015), and defined an interaction potential as the product of the ligand expression in the sender cell type and the expression of matching receptor in receiving cell type. This potential was computed separately for the NEC cells and the neonatal cells and the ratios of interaction potentials between NEC and neonatal samples were statistically assessed via random re-assignment of cells to the two groups (**Methods**). Consistent with increased activation of endothelial cells in our scRNAseq data, we found that NEC endothelial cells upregulated signaling to leukocytes via integrin receptors and cytokines including: macrophages via VCAM1-ITGB1 and IL6-F3, dendtritic cells via VCAM1-ITGB1/ITGB7, and IL6-IL6ST, and naïve T cells via MADCAM1-ITGB7, VCAM1-ITGB1/ITGB7/ITGA4 and IL6-IL6R/IL6ST (**Fig 6b-d**). Similarly, interactions with endothelilal cells through VCAM1-ITGB1/ITGB7/ITGA4 were upregulated in all T cell subsets including activated and regulatory T cells and MADCAM1-ITGB7 were additionally upregulated in Tregs (**Fig S7a**). NEC lymphatic endothelial cells, primarily interacted with leukocytes via cytokines and chemokines including: to macrophages via CCL3L1-CCR1, dendtritic cells via CXCL9/CXCL11-CXCR3 and CCL20-CCR6, and naïve T cells via CXCL9/CXCL10/CXCL11-CXCR3 (**Fig 6e-g**). Our ligand-receptor analysis indicated that NEC epithelial cells interact with macrophages through elevated levels of TNFSF9 (**Fig 6h**), and T cells via IL1B/IL1R1 and GZMB/PGRMC1 (**Fig 6i**). Fibroblasts interacted with macrophages through CXCL10-SDC4 and IL15-IL12RA/IL15RA (**Fig 6j)**. Finally, regulatory T cells interacted with macrophages through IL1B-IL1R1 while signaling to fibroblasts via TNFSF14-TNFRSF14 (**Fig S7b**). Numerous other interactions between cell types were identified (**Fig 6 and S7c-d**). Overall, epithelial-mesenchymal-endothelial-immune interaction were altered in NEC with increased ligand-receptor interactions between endothelial cells and immune cells; fibroblasts and immune cells, and epithelial cells and macropages and T cells and regulatory T cells.

## Discussion

The pathogenesis of NEC remains poorly understood and to date, few studies utilize single cell systems approaches to describe the altered interactions among various intestinal cell (Olaloye et al. 2021). Here, we present a single cell atlas with spatial resolution of the small intestine from neonates with and without NEC that identified 8 distinct cellular populations: macrophages, dendritic cells, B cells, T/NK cells, blood/lymphatic endothelial cells, fibroblasts, enterocytes and enteroendocrine cells. Our data exposes substantial changes in both abundances and transcriptional profiles of immune and non-immune populations in NEC SI.

Acute inflammation in intestinal tissues of patients with NEC was marked with inflammatory changes in numerous innate immune populations. We observed an increased abundance of inflammatory macrophages consistent with previous studies from our group and others (Emami et al. 2012; MohanKumar et al. 2016; Olaloye et al. 2021). Inflammatory macrophages were not only more abundant in NEC but also produced high amounts of inflammatory cytokines associated with recruitment of other immune cells to the sites of inflammation including IL1A and IL1B and TNFA (Orecchioni et al. 2019), with pathway analysis suggestive of inflammatory responses including TNFA signaling via NF*κ*B, cytokine receptor interactions, NOD like receptor pathways and TLR receptor signaling. NOD and TLR activation have been implicated in NEC pathogenesis both in human disease and in murine models, with the current study providing a context that Mϕ are responding to these signals (Afrazi et al. 2014; Hackam and Sodhi 2018; Cushing et al. 2017). Interaction analysis demonstrated that NEC-associated Mϕ interacted with both immune and non-immune cells within the SI. DCs were reduced in NEC mucosa consistent with our previous work (Olaloye et al. 2021). However, the DCs that were present in the SI exhibited a pro-inflammatory signature. Moreover neighborhood analysis of IMC data demonstrated an increase in DC: memory T cell interactions suggesting ongoing antigen presentation in NEC SI.

Previous work had implicated Th17 T cells in the pathogenesis of NEC demonstrating an influx of IL17 producing T cells that drive NEC inflammation in murine models (Jia et al. 2019; Pang et al. 2018b; Egan et al. 2016). We observed changes in T cell abundance associated with NEC with a reduction in naïve T cells and a trend towards an increase in activated and memory T cells. Although we did not observe IL17 upregulation in most cases, we did observe an upregulation of some genes classically involved in TH17 cells including BATF and CCL20 in NEC activated T cells (Maddur et al. 2012; Gaffen et al. 2014). This could perhaps be secondary to these samples being obtained later in the progression of NEC or suggest that IL17 independent pathways also contribute to NEC pathogenesis. We detected activated T cells with inflammatory signatures, indicative of increased cytotoxic activity in NEC SI. Surprisingly, major differences in TCRβ clonality, VDJ use, CDR3β length and presence of shared clones were evident in NEC compared to controls. Specifically, we noted an increase in the frequency of 3 in utero public clones. Although not previously observed in NEC, expansion of public clones has been described in infection and IBD (Werner et al. 2019; Musters et al. 2018). While the clones we identified are known to bind CMV antigens, it is unlikely related to the presence of active intestinal CMV infection in NEC cases as these clones were present in all NEC cases and found in majority of healthy fetuses (Stras et al. 2019). Public clones are known to be promiscuous (Dong et al. 2010) and these likely represent clones against bacterial or self-antigens that are similar to peptides found on CMV. Taken together, this suggests a pathogenic role and warrants further investigation into the nature of clonal expansion in NEC. Interestingly, our sequencing data showed shorter CDR3β length with fewer insertions and deletions compared to neonatal controls similar to what has been reported in IBD (Werner et al. 2019).

Regulatory T cells are critical for maintaining homeostasis and play a major role in intestinal inflammation through cell-cell interactions and secreted factors. Previous work on the role of Tregs in NEC had conflicting results with some demonstrating a reduction in Tregs in NEC (Pang et al. 2018a; Pang et al. 2018b; Weitkamp et al. 2009), while others suggested an increase in Tregs (Dingle et al. 2013). We did not observe significant abundance differences in Tregs between neonatal and NEC samples. Similar to a recent study demonstrating heterogeneity between NEC samples (Zhang et al. 2021), we observed large variability in the abundances of Tregs in NEC tissue as measured by scRNAseq, deconvolution of bulk RNAseq data and IMC. NEC Tregs showed increased CD62L and CCR7 expression, indicative of being recent thymic emigrants or natural Tregs rather than generated within the intestine (Wei et al. 2006). Furthermore, numerous genes that are classically associated with Treg identity or function including *CTLA4, ICOS, TIGIT*, and *TNFRSF4* were significantly down regulated, suggesting that NEC tregs are likely to have decreased supressive activity. Phosphorylation of p65(NFkB) has been previously shown to be important in maintaining Treg supresive signature (Oh et al. 2017). Consistent with the possibility that NEC Tregs have decreased suppressive activity, we found that p65 phosphorylation was decreased in NEC. Additionaly, NEC Tregs demonstrated an increase in STAT3 phosphorylations. STAT3 signaling has been shown to decrease FOXP3 expression and re-program Tregs towards the Th17 phenotype especially in the presence of IL6 and IL1β in the environment of secondary lymph organs (reviewed in (Colamatteo et al. 2019)).Both cytokines were elevated in the NEC tissues in our atlas. Although the regulation of Treg/Th17 balance is typically different in the intestinal mucosa, relying aditionally on retinoic acid and aryl hydroxylase receptor (AHR), similar IL6/IL1B reprogramming could potentially take place in the highly inflammed mucosa of NEC samples. IMC neighborhood analysis demonstrated overall reduced interactions of Tregs with other cell types including monocytes. Our ligand receptor interaction analysis of Tregs and other cell types highlighted upregulation of IL1β−IL1R interactions, previously shown to render Tregs less supressive (Raffin et al. 2013).

To determine non-immune signatures that could be leading to increased inflammation, we also investigated non-immune cell types found in the SI including endothelial cells, epithelial cells and fibroblasts and found reorganization and altered signaling in all three cell types. Enterocytes, particularly villus tip epithelial cells, were significantly reduced in NEC. A similar phenotype was observed in a “gut on a chip” model of NEC, where LPS treatement resulted in increased secretion of inflamatory cytokines including IL1β, IL6, CXCL8 and TNFa and a compromised intestinal barrier including villus blunting (Kim and Ingber 2013; De Fazio et al. 2021). Similarly, a recent study of human NEC samples demonstrated damage to the villus tips in patients with NEC, something we also observed in our study (Koike et al. 2020). The entorocytes present in NEC exhibited an inflammatory phenotype, with upregulation of numerous chemokines involved in recruiting immune cells with enriched ligand receptor interactions between epithelial cells and immune cells. Notably, upregulation of genes observed in NEC was similar to those previosuly identified in IBD including *REG1A, REG1B* and *DUOX2* (Puleston et al. 2005; MacFie et al. 2014; van Beelen Granlund et al. 2013; Kaemmerer et al. 2012). Furthermore, increased STAT3 phosphorylation in enterocytes was observed by IMC with upregulation of STAT3 dependent genes in the scRNAseq dataset, including *REG1A* (Mao et al. 2021), *REG1B, LCN2* (Ching et al. 2018) and *PLA2G2A*, antimicrobial peptides that are upregualted upon epithelial damage (Wittkopf et al. 2015).

Both fibrobalsts and endothelial cells were enriched in NEC SI and, similar to epithelial cells, demonstrated a proinflammatory program. Endothelial cells upregulated a number of genes involved in angiogenesis such as *CXCL1, CXCL8*, and *ALPN*, suggesting that some of the expansion in endothelial cells could be secondary to increased angiogenesis and endothelial cell proliferation (Russo et al. 2014). *CXCL1* and *CXCL8* are also involved in neutrophil recruitment, a known hallmark of NEC. Endothelial cell up-regulated genes were enriched in pathways involved in inflammation including TNFa, IL1β and IFN*γ* signaling and upregulated genes involved in leukocyte recruitment to sites of inflamamtion including those involved in rolling (in blood endothelium *SELE, SELP*) and adhesion (*ICAM1, VCAM1, MADCAM1*). Integrins interacting with VCAM1 and MADCAM1 include the *α*4β7(ITGA4/ITGB7) complex that is more specific for MADCAM1 and the *α*4β1 complex that preferentially binds to VCAM1. These integrins are upregulated on activated leukocytes, particularly lymphocytes, but have recently also been shown to be upregulated on innate cells, leading to leukocyte extravasation into intestinal high endothelial venules (Wang et al. 2010; Villablanca et al. 2014; Schippers et al. 2016; Zeissig et al. 2019). These receptor-integrin interactions were upregulated across the majority of T cell subtypes as well as dendritic cells and blood endothelial cells in NEC. Upregulation of *α*4β7 and *α*4β1 is thought to be pathogenic in inflamatory bowel disease (Zundler et al. 2017). Blockade of *α*4β7 with Vedolizumab, a humanized monoclonal antibody against *α*4β7, is effective for induction and maintenance of remission in IBD in multiple studies (Sandborn et al. 2019; Danese et al. 2019; Kopylov et al. 2019; Feagan et al. 2013; Sandborn et al. 2013). It is therefore feasible that a similar blockade could be effective in NEC.

In addition to the blood endothelium displaying a signature of intense activation, we observed altered expression of genes involved in decreased blood flow (*NOS3)*, increased vasocontriction (*EDN1)* and increased clotting *(F3, SERPINE1*, **Fig 4d**). Interestingly, while blood endothelial cells increased recruitment molecules that signal through integrins, lymphatic endothelial cells upregulated more cytokines and chemokines. In particular, CXCL9, CXCL10 and CXCL11 (Farnsworth et al. 2019) expression is atypical for such cell populations. These molecules are more associated with blood endothelial cells, where they regulate recruitment of T cells to the sites of inflammation. What role these chemokines might play in lymphatic endothelium is unknown. Overwhelming inflammation in endothelial cells may lead to dysfunction or death. Indeed, pathway analysis of DEG in NEC endothelial cells identified apoptosis as one of the upregulated pathways, a finding which we confirmed by TUNEL staining. This result potentially implicates blood/lymphatic endothelial cell death as contributing to intestinal inflammation. Fibroblasts demonstrated an increase in *IL1B* and *CSF3* and showed extensive interactions with other immune cells both by ligand-receptor interactions and by IMC neighborhood analysis. Previous work had identified inflammatory fibroblasts as potential drivers of IBD. It is intriguing to postulate that a similar mechanism might be at play in NEC (Kinchen et al. 2018).

NEC is a progressive disease and our atlas captured dysregulation in the small intestine in the subset of infants that required surgery. One limitation of our study is that it did not include infants who recovered from NEC with medical therapy. Validation that some of the markers identified here can also be detected in the blood of infants with NEC and their identification in infants with various stages of NEC would be interesting.

In summary, we provide a comprehensive atlas of immune-epithlial-endothelial-mesenchymal dysregulation in NEC accompanied by localization and ligand-receptor interaction analysis. Our study demonstrates profound inflammatory changes in NEC small intestine with increase in IL1β and TNFa producing macrophages, accompanied by inflammatory changes in endothelial, epithelial and fibrobasts cells, as well as increase in inflammatory signature in T cells with decreased suppressive signatures in Tregs. We also identify a number of potential interactions such as MADCAM1-*α*4β7 between endothelial and T cells that could represent future therapeutic targets for NEC treatment. Our data provides insights into the pathogenesis of NEC that will be instumental for future biomarker and therapeutic development.

## Supplementary Figure Legends

**Supplemental Figure 1: a**- t-stochastic neighborhood embedding (tSNE) of imaging mass cytometry data clustered by Phenograph color coded by cluster left-hand-side and condition (right-hand-side). **b**- Heatmap of normalized median epitope expression used to identify clusters.

**Supplemental Figure 2: a**- GSEA analysis of pathways enriched (red) or depleted (blue) in NEC samples compared to neonatal samples for dendritic cells (q value<0.2). **b**- t-stochastic neighborhood embedding of CD3^-^CD19^-^CD66b^-^ leukocytes from small intestine of infants with NEC (n=12) and neonates (neonatal n=4) adapted from (Olaloye et al. 2021) with percentages expressed in **Fig 2h**. Due to the influx and abundance of neutrophils in NEC affected mucosa, innate cell analysis of CyTOF data was performed on non-neutrophils (CD3^-^CD19^-^CD66b^-^)(Olaloye et al. 2021)

**Supplemental Figure 3: a-** Markers used to annotate the T cell clusters in Fig 3a-b. **b**- **-** Principal component analysis (PCA) plot of variable (V), - differential (D), and -joining (J) regions use in NEC and neonatal cases. **c**- Frequency of specific VDJ genes in neonatal and NEC samples. **d-** Complementarity-determining region 3 (CDR3) length, number of insertions and deletions in NEC and neonatal samples. **e**- Clonal representation of top clones in NEC and neonatal samples expressed as a proportion of the total clones in each sample. Black dots represents top three cytomegalovirus (CMV) clones, white dots are clones that were also notably present in small intestine of human fetuses as published in (Stras et al. 2019) and grey dots are other clones (see Methods and Supplementary **Table 11**). **f**. Differential gene expression between NEC and neonatal naïve T cells. Red dots are selected differentially expressed genes among the genes with q value<0.02 and fold change above 2 or below 1/2. Included are all genes with normalized expression above 10^−4^. **g** - Estimates of the proportions of ILCs and NK cells based on computational deconvolution of the sequencing data. Each dot is a sample, proportions were renormalized over all T cells, q-values are computed based on FDR correction for T cells subsets only. **h-i-** Differential gene expression between NEC and neonatal ILCs(h), NK cells (i). Red dots are selected differentially expressed genes among the genes with q value<0.02 and fold change above 2 or below 1/2. Included are all genes with normalized expression above 10^−4^.

**Supplemental Figure 4: a –** Expression of PDPN, CLDN5 in all cells in the atlas from Fig 1a-b. **b**-**c** - Expression of PDPN, CLDN5 (b) and CXCL10, CCL19, CCL21 (c) on blood and lymphatic endothelial cells only (Methods). **d-** Quantification of IF images denote tunel^+^Lyve1^+^ cells per high power field (hpf) (top) and area/intensity integration for Lyve-1 (bottom) in neonatal (n=4) and NEC (n=4) samples. p-values computed using Wilcoxon rank-sum tests.

**Supplemental Figure 5: a-b-** UMAPs colored by cell type of myeloid cells (a) and blood and lymphatic endothelial cells (b), **c -** expression of IL1B and TNF among myeloid cells (d) and SELE and ICAM1 in blood and lymphatic endothelial cells. **e**- Estimates of the proportions of distinct blood and lymphatic endothelial cell subsets, based on computational deconvolution of bulk sequencing data. Each dot is a sample, proportions were renormalized over all endothelial and lymphatic cells, q-values are computed based on FDR correction for endothelial and lymphatic cells only (Methods). MΦ− macrophages.

**Supplemental Figure 6 - a** – Re-clustered atlas of the fibroblast cluster. **b** – Cells annotated by condition. **c -** Differential gene expression between NEC and neonatal cells for the fibroblast cluster only. Included are all genes with normalized expression above 10^−4^. Red dots are the top 30 most differentially expressed genes among the genes with q value<0.01 and fold change above 2 or below 1/2.

**Supplemental Figure 7 – a-d -** Significantly elevated interactions between cell populations explored. Shown are the log-ratios between the interaction potentials in NEC and neonatal samples (Methods). Maps include the 25 significant interactions (q value <0.01) with highest fold change (Methods). In all interaction maps sender population is on the y axis, received is on the x axis.

## Supplementary Tables captions

**Supplementary Table1** - Markers of all cells. Markers for the eight cell type clusters, identified by the FindAllMarkers command in Seurat. Markers included have a log-fold above 1 and expressed in at least 50% of the cluster cells.

**Supplementary Table2** – IMC antibodies used for the experiments.

**Supplementary Table3** - DGE for all cell subsets. Differential gene expression between NEC and neonatal cells for the cell type subsets. Included are all genes with normalized expression above 10^−4^. p values were calculated using two-sided Wilcoxon rank-sum tests. q values were computed using the Benjamini-Hochberg false discovery rate correction.

**Supplementary Table4** – Amino acid sequences for clones identified in TCRβ analysis.

**Supplementary Table5** – IMC interactions **–** Adjacency interaction values in NEC and neonatal samples.

**Supplementary Table6** – Ligand receptor analysis table. Full table of ligand-receptor interaction analysis for all cell types.

**Supplementary Table7** – Samples used in the study. Demographics and allocation of samples used in the study.

**Supplementary Table8 –** Probe library sequences. Sequences for the smFISH probe libraries used in this study.

**Supplementary Table9** – Mix table for deconvolution. Bulk RNA sequencing count table of all 14 subjects (4 neonatal, 10 NEC) before correlation filtration in deconvolution analysis.

**Supplementary Table10** – Deconvolution results. Proportions table from cellanneal analysis.

**Supplementary Table11** - Signature table for deconvolution. Mean expression of major cell types from single cell data. Data in sum-normalized.

## Methods

### Intestinal tissue acquisition and storage

Fresh small intestinal (SI) tissue from human neonatal and NEC samples were obtained from surgical resections in infants with IRB approval (**Supplementary Table 7**). No consent was obtained for neonatal samples as they were collected without any identifying information under a discarded specimen protocol that was deemed non-human research by the University of Pittsburgh IRB (IRB# PRO17070226). For single cell sequencing and paraffin blocks, tissue was cryopreserved (Stras et al. 2019). Briefly, intestinal tissue samples were cut into sub-centimeter pieces and cryopreserved in freezing media (10% dimethyl sulfoxide (DMSO) in fetal bovine serum (FBS)) in a slow cooling container (Mr. Frosty) at -80°C for 24 hours, then transferred into liquid nitrogen for long term storage. For paraffin blocks, tissue was fixed in 4% formalin for 48 hrs, transferred to ethanol until embedded in paraffin. Blocks were stored in paraffin until sectioned for analysis.

### Intestinal Tissue Digestion

Cryopreserved samples were processed as previously described (Konnikova et al. 2018). Briefly, intestinal tissue samples were quickly thawed and washed in RPMI medium plus 10% FBS (Corning), 1X GlutaMax, 10mM HEPES, 1X MEM NEAA, 1 mM sodium pyruvate, 100 I.U/mL penicillin and 100 μg/ML streptomycin. Next, intestinal tissue was incubated overnight in the same media with 1 μg/mL DNase 1 and 100 μg/mL collagenase A. Tissue dissociation was performed on the gentleMACS Octo Dissociator with heaters (Miltenyi Biotec) using the heated human tumor protocol 1. Tissue was then filtered through a 70- μm nylon mesh cell strainer (Sigma) to make a single cell suspension.

### Single cell RNA sequencing

Single cell suspension obtained from SI tissue digestion was washed in T-Cell Media twice followed by enrichment for live cells with the dead-cell removal kit using the MACS Cell separation system (Miltenyi kit # 130-090-101). Viable cells were counted with trypan blue using a hemocytometer. Using the MS or LS columns depending on the number of cells obtained, cells were separated into a CD45^+^ and CD45^-^ fractions using the MACS Cells separation system with the CD45 bead enrichment (Miltenyi kit # 130-045-801). Cells were again counted on a hemocytometer with trypan blue. Libraries were made using the Chromium Next GEM Single Cell 3’ kit v3.0 (10X Genomics) with a target of 5,000 cells. 3’ GEM libraries were made separately, once from the CD45^-^ fraction and once from the entire single cell suspension from the tissue digest without any selection. 3’ GEM libraries were sequenced at Medgenome. Sequencing was performed on HighSeq lane with two samples per lane (Xin et al. 2020; Luo et al. 2020) or on an S4 NovaSeq lane.

### Bulk sequencing

RNA was extracted from snap frozen intestinal tissues samples using the Qiagen All 422Prep Kit (#80204). cDNA synthesis was prepped by the Yale Genomics core. Quality 424control was completed by MedGenome via Qubit Fluorometric Quantitation and TapeStation 425BioAnalyzer. Libraries were sequenced on the NovaSeq6000 for Paired End 150 base pairs for 42690 million reads per sample. Paired-end reads fastq files were mapped with Hisat2.

### CyTOF staining and analysis

Sample were stained with antibody cocktail per previously published protocol (Stras et al. 2019) and incubated with 191Ir/193Ir DNA intercalator (Fluidigm) and shipped overnight to the Longwood Medical Area CyTOF Core. Data was normalized and exported as FCS files, downloaded, and uploaded to Premium Cytobank® platform. Gating and analysis with cytofkit (Chen et al. 2016) as published (Stras et al. 2019). Cluster abundance was extracted, analyzed and plots generated using R. In addition, mean expression values for neutrophil clusters was extracted for antibodies of interest were analyzed in Prism 9.

### IMC staining and analysis

Biopsy-sized pieces of small intestinal tissue were fixed in formalin on day of collection and subsequently embedded in paraffin in batches. Formalin-fixed, paraffin embedded (FFPE) tissue was sectioned into 4-5 μM thick sections. Slides were deparaffinized using xylene and alcohol and placed in 1X antigen retrieval buffer (R&D, #CTS013) at 95°C for 20 minutes. Next, slides were washed in distilled H20 (ddH20) and Dulbeco’s Phosphate Buffered Solution (DPBS, Gibco). Tissue was blocked with 3% BSA in DPBS for 45 minutes at room temperature. Overnight incubation of antibodies (**Supplementary Table2**) diluted in 0.5% BSA in DPBS at 4°C was performed. Slides were rinsed in DPBX with 0.1% TritonX100 twice and DPBS twice. Counterstain was performed with 100 μM LipoR-Ln115 and Ir-intercalator (1:2000) in ddH20 at room temperature for 30 minutes. Slides were rinsed in ddH20 and then air dried.

### Selection of areas of interest

Regions of small intestinal tissue were selected manually to capture all layers of the intestine. The same sized area was scanned in all samples.

### IMC image acquisition

The Helios time-of-flight mass cytometer (CyTOF) coupled to a Hyperion Imaging System (Fluidigm) was used to acquire data. An optical image of each slide was acquired using Hyperion software and areas to ablate were selected as described above. Laser ablation was performed at a resolution of 4 μM and a frequency of 200 Hz. Data from slide was acquired over 2 consecutive days in total of 18 image stacks from 9 samples.

### IMC data segmentation and analysis

Data from Hyperion extracted as MCD and .txt files that were visualized using Histocat++ (Fluidigm). Further analysis of image data was performed using a recently published IMC segmentation pipeline (Damond et al. 2019) that was adapted to our dataset. Briefly, a Python script (https://github.com/BodenmillerGroup/imctools) was used to convert text format (.txt) files from data acquisition to tiff images. Spillover compensation was performed to minimize crosstalk between channels. The images where segmented in two steps. First ilastik (Berg et al. 2019), an interactive machine learning software was used to classify pixels as nucleus, cytoplasm or background components. Training of the Random Forest Classifier was performed on 125 × 125-pixel sub stacks generated from original images using relevant markers (e.g CD45, CD3, CD14, CD163, panCK, SMA). CellProfiler was used to identify nuclei, define cell borders and generate cell masks and identify single cell data from original images. Files and masks were loaded to histoCAT 1.7.6.1 (Schapiro et al. 2017), visualized and analyzed as follows. Images from Figure 5 were generated using the image visualization option in HIstocat 1.7.6.1 (Schapiro et al. 2017). Phenograph clustering of cells was performed using major lineage markers. IMC raw counts and median metal intensity for each cell for phosphor markers were extracted as .csv for each cluster and data graphed using Graphpad® Prism 9. The sum of cells in each cluster was extracted from .csv files and used to calculate relative abundances. Individual expression data from each cell was exported for markers of interest and graphed using box and whisker plots in Graphpad® Prism 9.

Nearest neighborhood analysis was performed in histoCAT identifying neighboring cells within 4 pixels (1 pixel ∼ 1 micron) to identify with interactions present in >10% of images with a of p-value of 0.05 and 999 permutations. The program returns a matrix of cell-cell interactions that occur more frequently than random chance with the frequency of interactions expressed as a fraction of total images. There were 6 images for neonatal (2-400×400 microns sections scanned for each of the 3 samples) and 10 images for NEC (2-400×400 microns sections scanned for each of the5samples). Interactions within a row answer the question: is cell type × in the neighborhood of cell type 7 (or is Y surrounded by X). Those within a column answer the question: is cell type 7 in the neighborhood of cell type × (x surrounded by 7). Interaction adjacency values are presented in **Supplementary Table 5** . Dot plots (Figure 6a) were produced by removing all adjacency interaction values involving the ‘other’ cluster and sorting the differences between the interaction values in NEC and neonatal samples. The figure showed the top 20 adjacency interactions among the interactions with positive values in NEC, sorted from largest to smallest difference between NEC and neonatal samples (red dots) and the bottom 20 interactions with negative values in NEC, sorted from the smallest to the largest difference between NEC and neonatal samples (blue dots).

### TUNEL

Paraffin embedded human intestinal tissue samples were sectioned, deparaffinized with xylenes (Sigma 534056-4L) and ethanol (Fisher BP2818-4) and fixed in 4% paraformaldehyde (PFA). Using the Click-iT™ Plus TUNEL Assay (Invitrogen C10619), samples were then permeabilized with Proteinase K solution. The TdT reaction was performed on the samples for 60 minutes at 37°C. Lastly, the Click-iT™ Plus reaction was performed on the samples for 30 minutes at 37°C.

### Immunofluorescence

Samples were blocked with 10% horse serum and incubated with primary antibodies (LYVE-1, 1:19, Biotechne AF2089). Samples were then incubated with secondary antibody (1:750, Invitrogen, A3216). Images were taken at 20x using Echo® Revolve microscope.

### Quantification

Fluorescence was quantified using ImageJ/Fiji. Equal size areas on immunofluorescent images were measured for integrated density and mean gray value. Mean gray values of background regions without fluorescence were also measured. Corrected total cell fluorescence (CTCF) was calculated from integrated density, area, and background fluorescent values.

### smFISH

smFISH was performed on frozen sections, with a modified smFISH protocol that was optimized for human intestinal tissues based on a protocol by Massalha, H. *et al*. (Massalha et al. 2020). Intestinal tissues were fixed in 4% Formaldehyde (FA, J.T. Baker, JT2106) in PBS for 1-2 hour and subsequently agitated in 30% sucrose, 4% FA in PBS overnight at 4°C. Fixed tissues were embedded in OCT (Scigen, 4586). 6-8µm thick sections of fixed intestinal tissues were sectioned onto poly L-lysine coated coverslips and fixed again in 4% FA in PBS for 15 minutes followed by 70% Ethanol dehydration for 2h in 4°C. Notably, unlike Massalha *et al*., tissues were not permeabilized with PK prior to hybridization.

Tissues were rinsed with 2× SSC (Ambion AM9765). Tissues were incubated in wash buffer (20% Formamide Ambion AM9342, 2× SSC) for 30-60 minutes and mounted with the hybridization mix. Hybridization mix contained hybridization buffer (10% Dextran sulfate Sigma D8906, 30% Formamide, 1□mg/ml E.coli tRNA Sigma R1753, 2× SSC, 0.02% BSA Ambion AM2616, 2□mM Vanadyl-ribonucleoside complex NEB S1402S) mixed with 1:3000 dilution of probes. Hybridization mix was incubated with tissues for overnight in a 30°C incubator. SmFISH probe libraries (**Supplementary Table8**) were coupled to Cy5, TMR or Alexa594. After the hybridization, tissues were washed with wash buffer for 15 minutes in 30°C, then incubated with wash buffer containing 50□ng/ml DAPI (Sigma, D9542) for 15□minutes in 30°C.

Tissues were transferred to GLOX buffer (0.4% Glucose, 1% Tris, and 10% SSC) until use. Probe libraries were designed using the Stellaris FISH Probe Designer Software (Biosearch Technologies, Inc., Petaluma, CA), see **Supplementary Table 8**.

### Imaging

smFISH imaging was performed on Nikon eclipse Ti2 inverted fluorescence microscopes equipped with 100x and 60x oil-immersion objectives and a Photometrics Prime 95B using the NIS element software AR 5.11.01. Image stacks were collected with a z spacing of 0.3µm. Identification of positive cells was done using Fiji (Schindelin et al. 2012).

### SELE quantifications

SELE positive cell was defined as positive with more than two smFISH dots. Cells were manually quantified and p-value was calculated using two-sided Wilcoxon rank-sum test. Two subjects for each condition (NEC/ neonatal) were quantifies over all intestinal layers (villi, crypts, submucosal area, muscle layers and serosa). Each dot represents the number of SELE+ cells in one imaging field.

### Computational analysis

#### Single cell analysis

The single-cell RNAseq data was processed using Cell Ranger 4.0.0 pipeline to align reads and generate a count matrix. Cell-free RNA, which often creates a background in scRNAseq was removed as follows: background cells were defined as cells with 100-300 UMIs and mitochondrial fraction below 50%. The average UMI counts for each gene in the background cells was subtracted from the respective counts in all other cells. Subsequently, cells with less than 200 expressed genes were filtered out and genes that were expressed in less than three cells were removed from data. Single cell data was analyzed using R software version 4.0.2 and Seurat package (version 3.2.2(Stuart et al. 2019)).

All subjects were merged and cells with less than 1900 UMIs, less than1000 genes per cell or mitochondrial fraction above 30% were filtered out. In addition, cells with a fraction of erythrocyte markers (“HBG2”, “HBA2”, “HBA1”, “HBB”, “HBG1”, “HBM”, “AHSP”, “HBZ”) above 10% were filtered out. Data was normalized and scaled using the SCTransform function, with regression of sum of UMIs (vars.to.regress = “nCount_RNA”). PCA was calculated based on the variable genes with the exception of mitochondrial (“^MT-”) and ribosomal (“RP[LS]”) genes. These genes were manually removed from the PCA, since they are prone to batch-related expression variability. Clusters were annotated based on markers from literature (see **Supplementary Table 1** for a complete list of markers obtained using FindAllMarkers in Seurat).

Cell type structures (Macrophages and dendritic cells, T cells, endothelial and lymphatic endothelial cells, enterocytes and fibroblasts) were computationally extracted from the complete atlas based the cluster annotations. Cells were normalized and scaled using the SCTransform function, with regression of sum of UMIs (vars.to.regress = “nCount_RNA”). PCA was calculated based on the variable genes with the exception of mitochondrial (“^MT-”) and ribosomal (“RP[LS]”) genes. Within the T cells, The Tregs+TRMs cluster was subset and re-clustered, to enable proper splitting of the two cell populations. The resulting split cell annotations were presented in the main T cell UMAP. Re-clustering of Fibroblasts subset revealed two clusters-fibroblasts and neurons.

DGE analysis used two-sided Wilcoxon rank-sum tests. DGE was only performed for genes with sum-normalized expression above 10^−4^ (or above 5×10^−5^ in crypt-mid-bottom enterocytes). In addition, only genes that were expressed in 2 or more subjects and in at least 5 cells in each of the two subjects were retained. Q-values were computed using the Benjamini-Hochberg false discovery rate correction (Benjamini and Hochberg 1995). Gene Set Enrichment Analysis (GSEA) included genes with sum-normalized expression above 10^−4^. The rnk file for GSEA included the sorted log2-fold expression between NEC and neonatal samples. In the T cell DGE TRMs were not shown since neonatal cells were underrepresented.

#### Ligand receptor analysis

Ligand receptor analysis was performed following Martin et al. (Martin et al. 2019). Briefly, this analysis computes for each pair of sender cell type A and receiver cell type B, and for each pair of ligand L and its matching receptor R, an interaction potential. The interaction potential is defined as the product of the mean expression of ligand L in cell A and the mean expression of receptor R in cell B. The interaction potentials are computed separately for the NEC cells and the neonatal cells, and their ratio is compared against a randomized dataset. In the randomized dataset, cells were re-assigned to the NEC and neonatal identities at random within each cell subtype, thus preserving the total number of NEC and neonatal cells. Random re-assignment was repeated 100 times and a probability that the observed ratio is higher than that expected by chance was computed using the normal distribution over the standardized ratio. The standardized ratio was defined as the difference between the real ratio and the average randomized ratio divided by the standard deviation among all randomized ratios. We only considered interactions that obeyed the following criteria: both ligand and receptor expressed in 10 or more cells in each of the respective cell types and in at least 10% of the cells in the cell type cluster in NEC or neonatal. Q-values were computed using the Benjamini-Hochberg false discovery rate correction (Benjamini and Hochberg 1995). For each pair of cell types of interest, the top 25 interactions with q-value <0.01 (for some pairs less than 25 interactions passed this criteria) sorted by ratio were selected and plotted (**Supplementary Table 6**).

#### Deconvolution analysis

Bulk data was filtered to include only coding genes, taken from Human_GRch38_91_ensemblBioMart and normalized to the sum of reads for each sample (**Supplementary Table 9**). Computational deconvolution was performed with cellanneal (Buchauer and Itzkovitz 2021), (**Supplementary Table 10**). Only samples with Spearman correlations above 0.3 between the mixture data and the synthetic mixtures were considered. For the signature file we included the mean expression per cell type for the original 8 cell type clusters shown in **Figure 1a** or for all cells, after breaking up cell-type clusters into their sub-types (**Supplementary Table 11**). Since the two subsets of inflammatory macrophages showed a clear separation by disease status they were grouped together. For deconvolution of cells broken into sub-types, estimated proportions for each cell type were normalized internally over the respective cell subtype. P-values were calculated using two-sided Wilcoxon rank-sum tests. Q-values were computed using the Benjamini-Hochberg false discovery rate correction (Benjamini and Hochberg 1995) and were calculated over the internally normalized proportions of each respective cell sub-types.

#### DNA Extraction

DNA was extracted from snap frozen intestinal tissue samples with the Wizard® Genomic DNA Purification Kit (Promega) per manufacturer’s instructions for animal tissue.

#### TCRβ Repertoire Library generation and analysis

Primers for various V and J gene segments in the *TRB* loci were used for amplification of rearranged CDR3B for each genomic DNA sample (ImmunoSeq TRB Survey Service, Adaptive Biotechnologies, Seattle WA, USA). Survey level of up to 500, 000 reads/sample was used. Libraries were purified, pooled and subjected to HTS using Illumina technology (Illumina Inc., San Diego, CA, USA) per manufacturer’s protocol. ImmunoSeq software online was used to analyze clonality, percentage of T or B cells, sample overlap, percent productive template, CDR3B length and VDJ use. Graphical representation of each repertoire is represented with hierarchical tree maps using available software (www.treemap.com).

## Data Availability

All data generated in this study is available at the Zenodo repository under the following https://doi.org/10.5281/zenodo.5813398.

## Author’s Contribution

S.I, L.K., O.O. and A.E. conceived the study. O.O was involved in sample collection, processing and preparation for single cell analysis. B.M. was involved in bulk RNAseq preparation. X.A., F.W. and K.C. were involved in library preparation. A.E. performed computational analysis and smFISH experiments. O.O performed suspension and imaging mass cytometry experiments and image analysis. T.S. performed the TUNEL and IF staining as well as image quantification. L.W and D.S performed NGS analysis. J.S.P. and R.W.P. assisted with endothelial cell analysis. L.K. and S.I. supervised the entirety of the project. All authors approved the manuscript.

