## Supplementary figures and images for "Single cell atlas of the neonatal small intestine with necrotizing enterocolitis"

### supl Fig 1

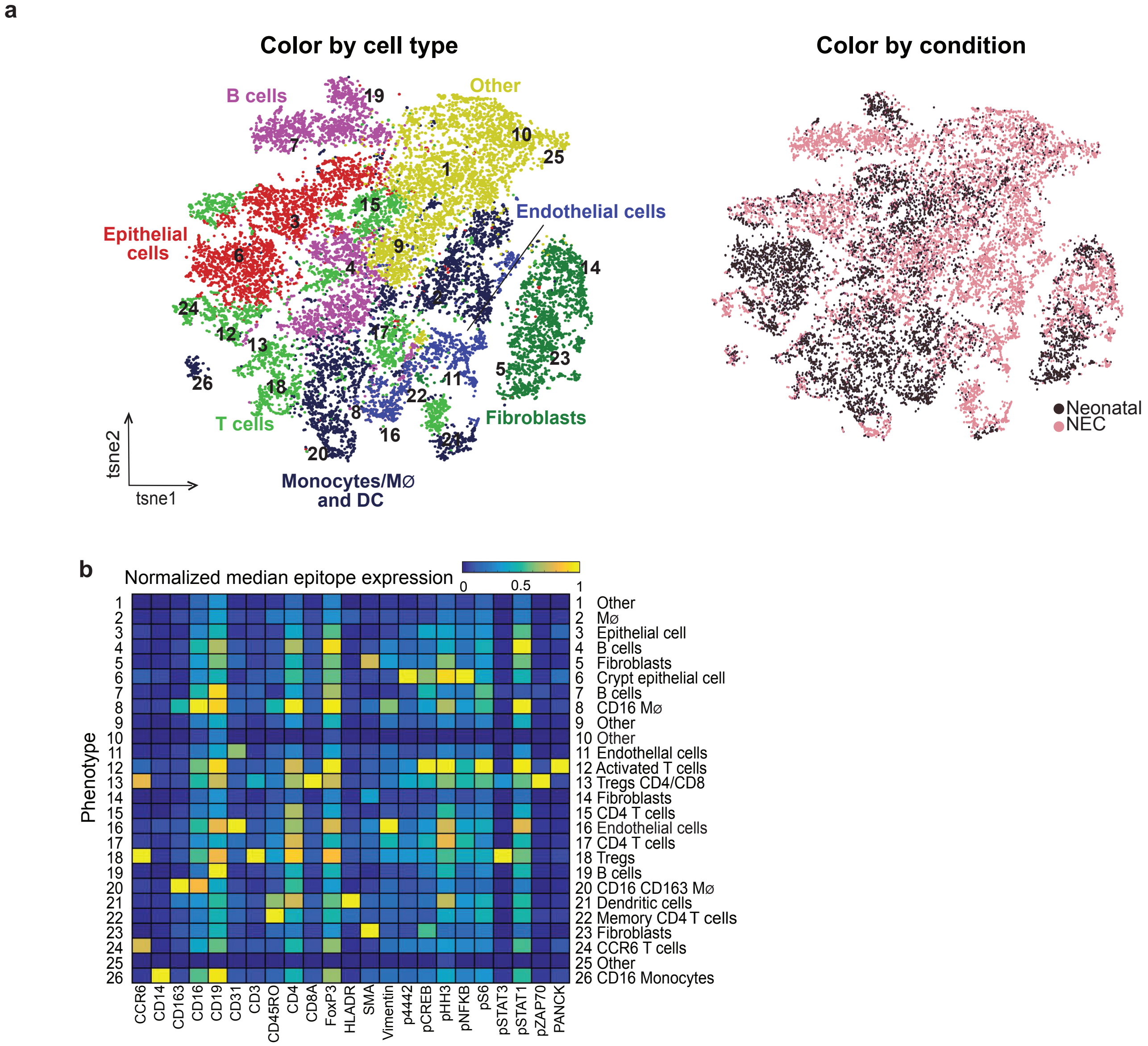

### supl fig 2

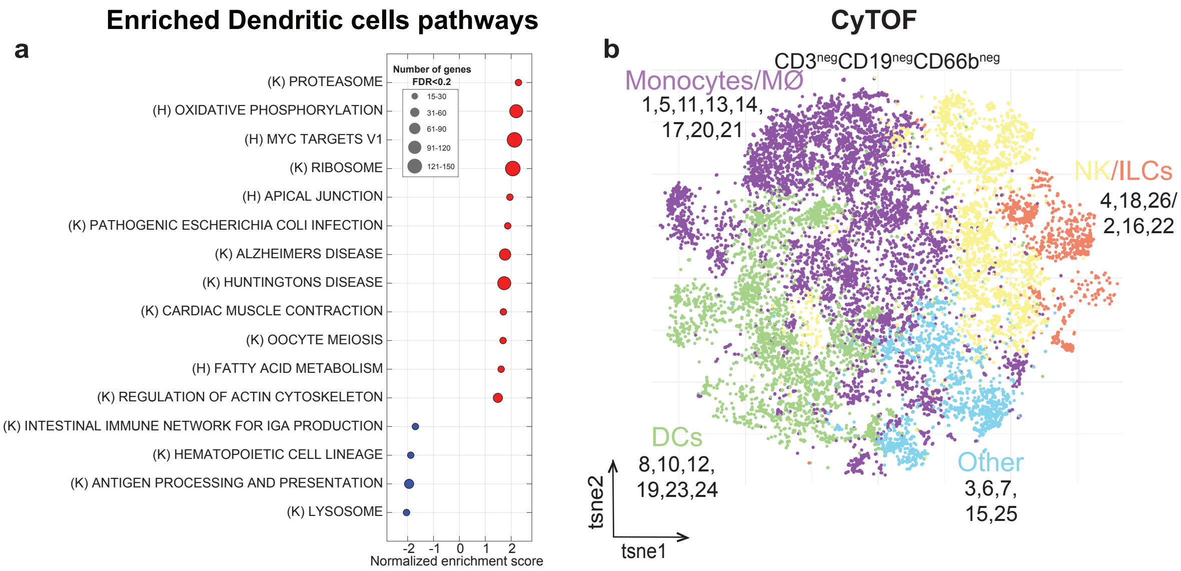

### supl fig 3

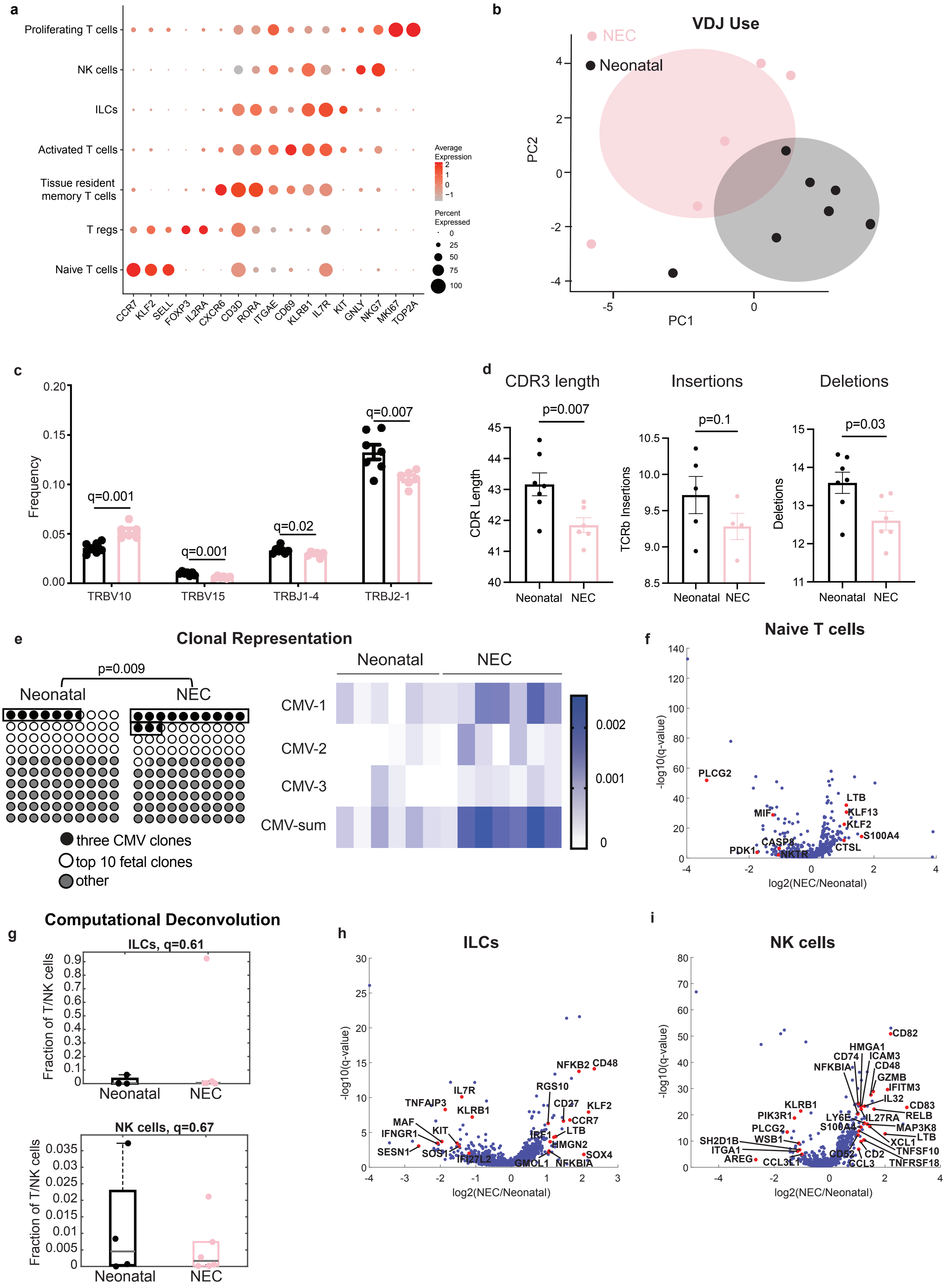

### supl fig 4

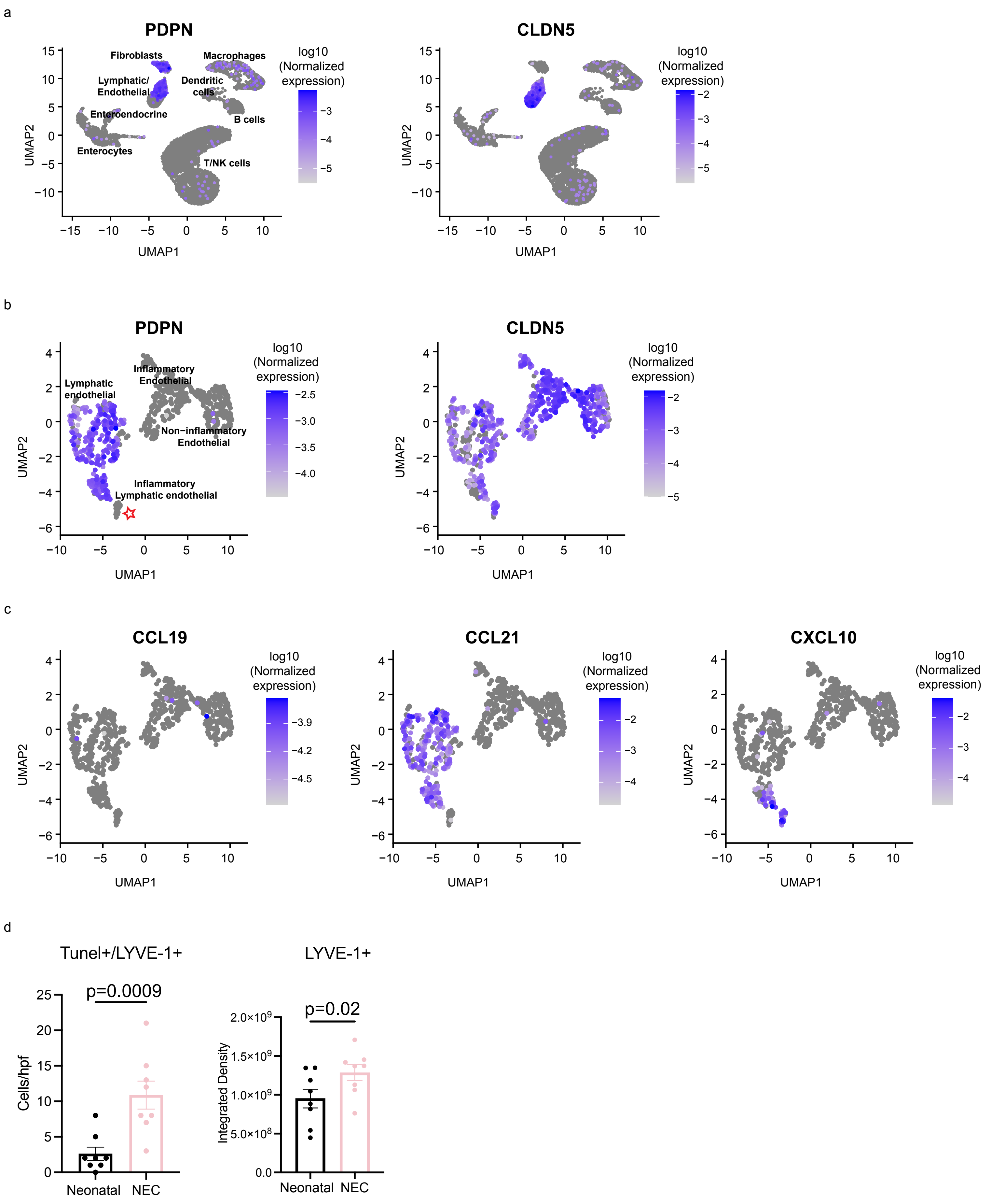

### supl fig 5

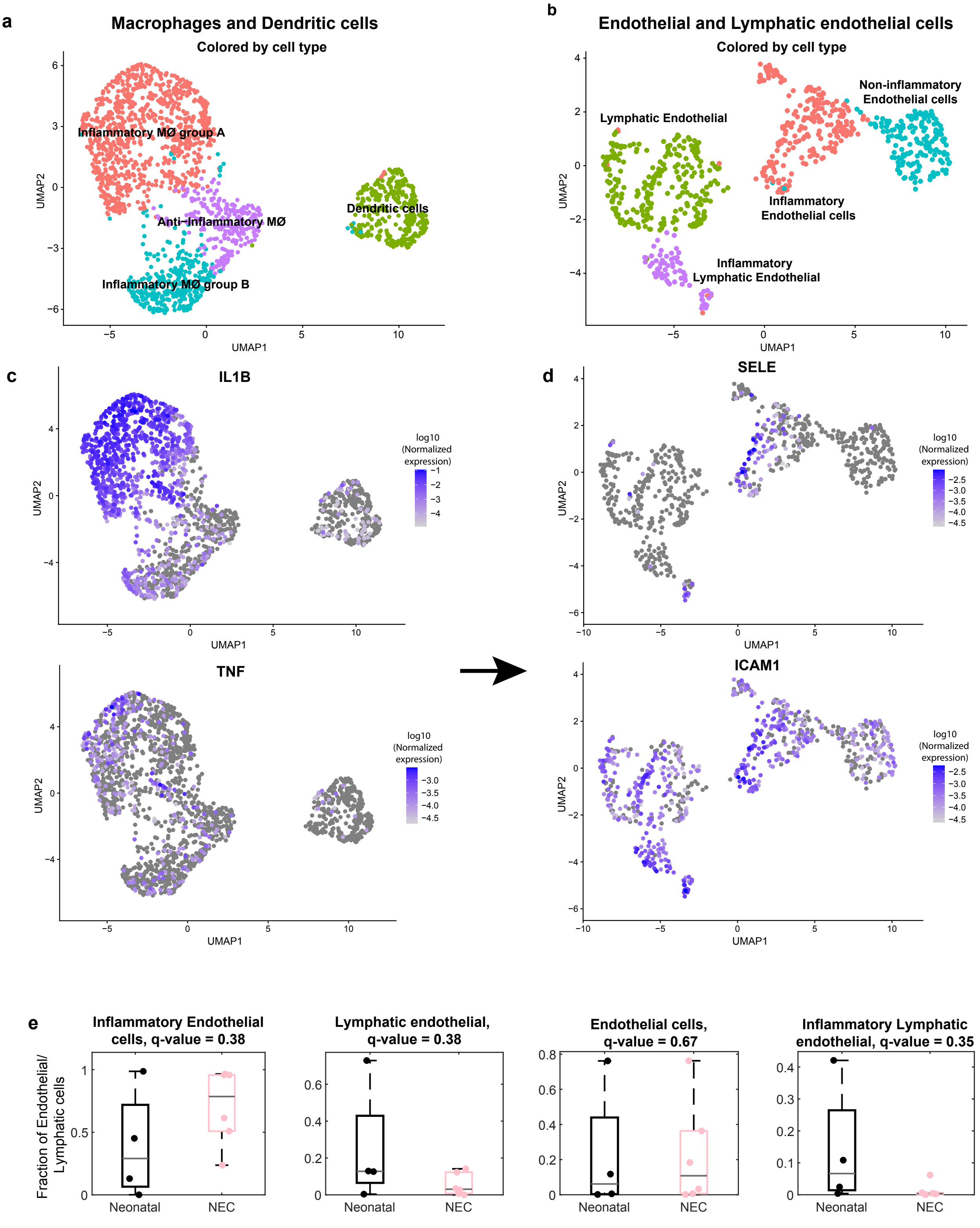

### supl fig 6

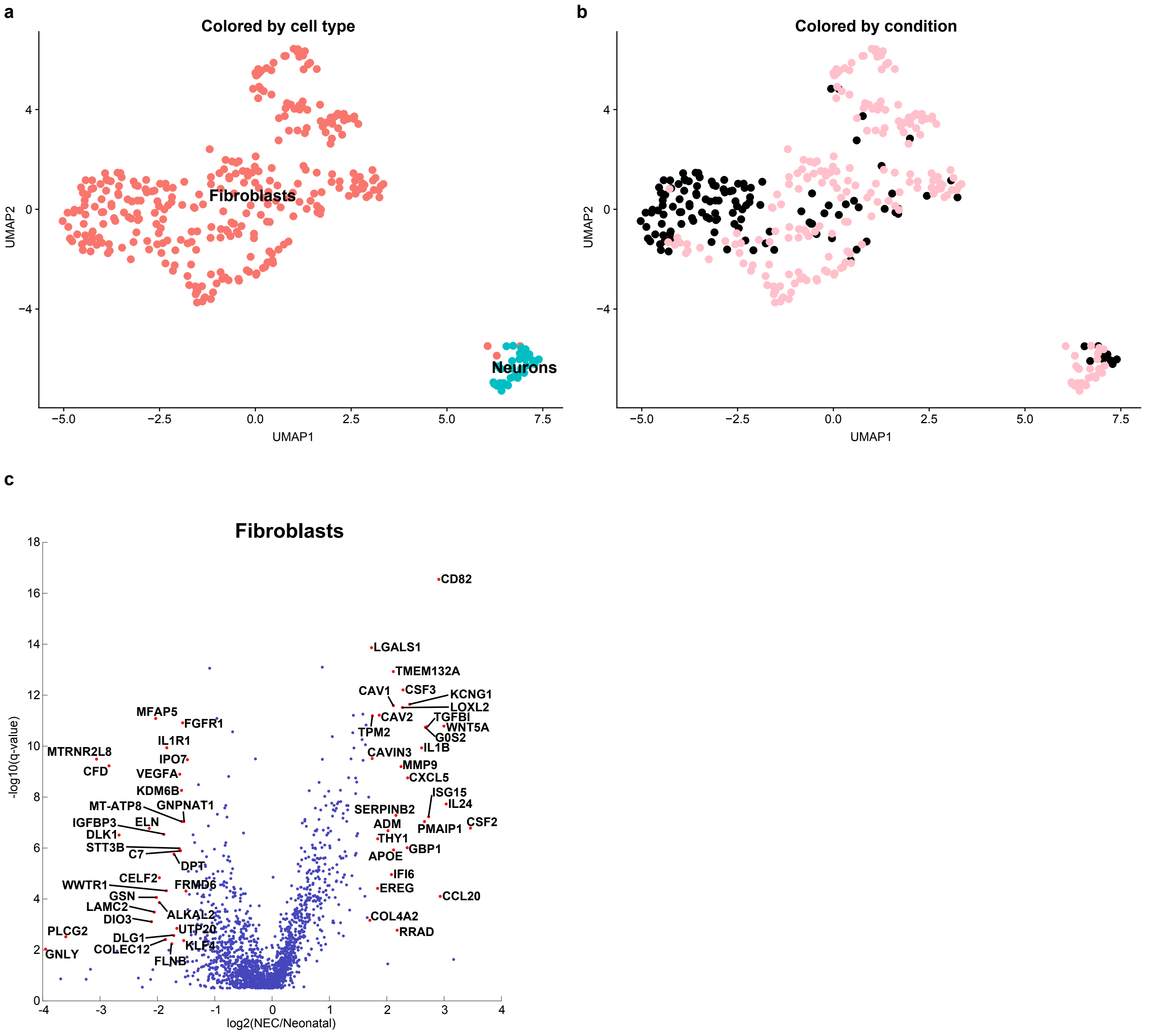

### supl fig 7

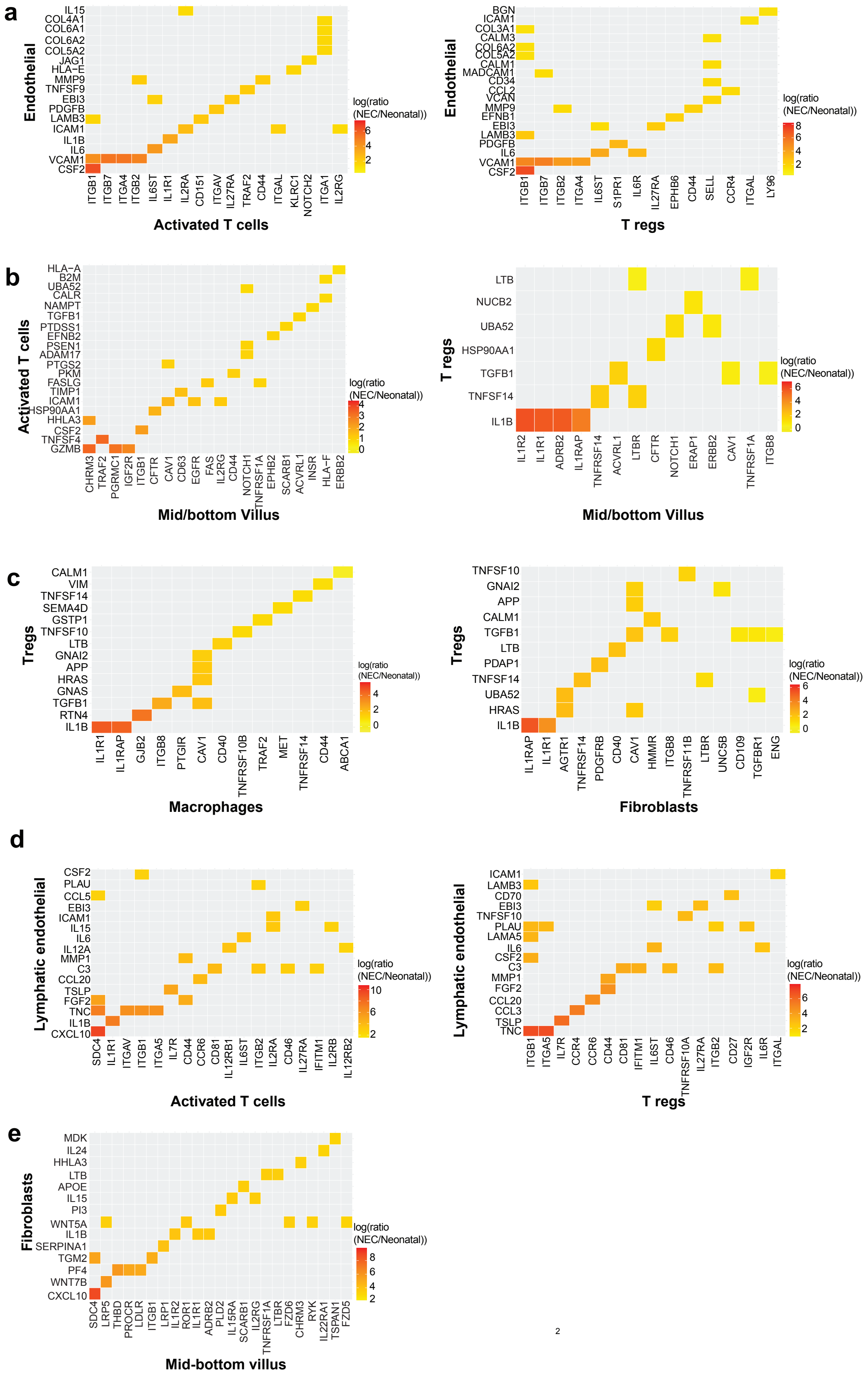
